# Cross-species meta-analysis reveals determinants of homing gene drive performance

**DOI:** 10.64898/2026.02.28.708699

**Authors:** Sebald A. N. Verkuijl, Edward R. Ivimey-Cook, Bairu Liu, Michael B. Bonsall, Philip T. Leftwich, Nikolai Windbichler

**Author notes:** These authors jointly supervised this work (Michael B. Bonsall, Philip T. Leftwich, Nikolai Windbichler).

## Abstract

Homing gene drives can bias their inheritance above Mendelian expectations, but reported outcomes vary widely. We compiled a cross-species dataset of nearly one million scored progeny from 42 publications reporting CRISPR/Cas9 endonuclease-based gene drives in 10 model, pest, and vector species. Using multilevel meta-analytic models, we evaluate biological, cross, and transgene design factors as predictors of biased drive inheritance. Species is the strongest predictor, but most heterogeneity remains unexplained; design features each explain a modest fraction of the remaining variation, with large construct-to-construct differences pointing to the full combination of design choices rather than any single factor. Nuclease expression timing, the most common optimization target, has limited predictive value after accounting for correlated factors, and predictions do not transfer well between species. Maternal nuclease deposition has a marginal effect on drive inheritance but dramatically increases somatic phenotype rates in offspring, revealing a tissue-of-action rather than repair-outcome effect. An interactive web tool enables community analysis of this dataset, which will guide the design of more efficient gene drives for genetic vector control, invasive species management and other applications. https://sverkuijl.shinyapps.io/GeneDrive/

## 1 Introduction

Gene drives are genetic systems that bias their own inheritance above the Mendelian expectation of 50%, enabling a selected allele to spread through a population even when it confers no fitness advantage ^1^. Among proposed mechanisms, homing gene drives are the most widely studied. These drives use a site-specific endonuclease to cut the homologous chromosome in heterozygous germline cells; the DNA double-strand break is then repaired by homology-directed repair (HDR) using the drive-carrying chromosome as template, converting the cell from heterozygous to homozygous for the drive allele ^2^. If this conversion occurs in the germline, the drive allele is transmitted to a majority of the progeny. Synthetic homing drives have potential applications in vector-borne disease control, agricultural pest management, and conservation ^1^.

Early work using meganucleases and other programmable nucleases in *Anopheles gambiae* and *Drosophila melanogaster* reported substantial inheritance bias, suggesting that efficient gene drives based on the homing mechanism could be readily engineered in the germline of metazoans ^3–6^. The development of flexible CRISPR-Cas9 as a programmable nuclease platform ^7^ greatly accelerated homing drive research; the first CRISPR-based homing drives reported strong inheritance bias in *Dr. melanogaster* ^8^ and *Anopheles* ^9;10^, prompting rapid testing across a widening range of species and target genes. However, this also revealed wide performance variation, and it became clear that the high rates of drive achieved in *Anopheles* mosquitoes were not readily generalisable.

Much of the ensuing optimisation effort has focused on the developmental timing of nuclease expression, on the premise that restricting Cas9 activity to meiotic cells should favour HDR over competing repair pathways, typically by testing panels of germline-specific promoters ^11–21^. Yet in practice, multiple design parameters for example promoter choice, gRNA target, transgene insertion sites, and genetic background differ widely between implementations, and reporting standards vary across laboratories. It has therefore been difficult to attribute performance differences to individual factors from the published literature alone.

No systematic, quantitative synthesis of homing-drive data exists. Here, we compile a cross-species dataset of nearly one million individually scored progeny from 42 publications spanning 10 species, and apply multilevel meta-analytic models to evaluate the contributions of biological, transgene design, and experimental factors to drive inheritance. We also provide an interactive web tool for the research community to perform customised analyses and inspect individual crosses in the underlying dataset.

## 2 Results

### 2.1 Dataset compilation

We retrieved 1,836 unique publication records from Pubmed and the Web of Science. Following the screening process and the application of exclusion criteria, we extracted cross-level data from 42 publications (Fig S1, Supplementary Data S1). We excluded studies in cell culture and single-celled organisms, and restricted *Drosophila melanogaster* to publications testing promoters other than the repeatedly used vasa and nanos elements alone, thus focusing on optimisation strategies comparable to those in pest species. Despite this restriction, data from *Dr. melanogaster* constitutes the second-largest set of publications. Non-CRISPR homing drives (principally I-*Sce*I) were not included because inheritance rates were generally not reported in a format compatible with a common database ^3–5^.

The resulting dataset comprises nearly one million individually scored F2 progeny across 10 species, with nearly 100 primary metadata fields recorded per cross (Supplementary Data S2), including over ten secondary outcomes beyond drive allele inheritance. Data extraction was broadly inclusive, with additional filters applied specifically for drive inheritance shown in Fig S2.

**Table 1:** Species distribution in the manuscript corpus. F2 progeny counts for the full dataset and each primary analysis population (drive allele inheritance and nuclease-induced somatic phenotype). See **Supplementary Data S2** for full dataset characteristics.

| Species | Pubs* | Crosses | F2 Total | F2 Drive | F2 Somatic |
| --- | --- | --- | --- | --- | --- |
| *Ae. aegypti* | 6<sup>19;21–25</sup> | 193 | 268,298 | 223,673 | 71,193 |
| *An. gambiae* | 16<sup>10;14;20;26–38</sup> | 193 | 231,881 | 108,539 | 60,898 |
| *Dr. melanogaster* | 7<sup>11;12;15–17;39;40</sup> | 169 | 138,080 | 125,323 | 38,469 |
| *P. xylostella* | 1<sup>41</sup> | 118 | 129,917 | 125,337 | 66,056 |
| *An. stephensi* | 5<sup>9;42–45</sup> | 101 | 99,631 | 89,990 | 50,382 |
| *Dr. suzukii* | 2<sup>17;46</sup> | 25 | 36,950 | 36,879 | 0 |
| *Ce. capitata* | 1<sup>18</sup> | 15 | 36,206 | 36,206 | 24,374 |
| *An. coluzzii* | 2<sup>13;28</sup> | 29 | 30,803 | 27,485 | 9,030 |
| *Cx. quinquefasciatus* | 1<sup>47</sup> | 7 | 6,196 | 6,196 | 2,063 |
| *M. musculus* | 3<sup>48–50</sup> | 14 | 2,545 | 2,544 | 0 |
| **Total** | **42** | **864** | **980,507** | **782,172** | **322,465** |

\* Multi-species papers contribute to multiple rows; per-species counts exceed the total.

### 2.2 Species is a strong, but incomplete, predictor of drive inheritance

Drive inheritance data by publication are shown in Fig S3. The dataset spans 2015 to mid-2025; reported inheritance rates have not systematically improved over time for any species (Fig S4; meta-regression slope 0.0201 logit units per year, SE = 0.0660, *p* = 0.7612; corresponding to a predicted change of +1.8% at a 75% baseline over five years). The % change per logit unit at each reference point is shown in Fig S5. Figure-by-figure statistical reports are listed in Supplementary Data S3 and S4.

Where data are reported per parent, parent-to-parent variability in inheritance rates was greater than expected from random sampling alone in crosses where drive activity is expected (Fig S6). Given limited per-parent data and in-consistent reporting conventions, we analysed pooled cross-level data throughout. For visual clarity, replicate crosses defined as such within publications are plotted as single data points (Fig S7).

Unless otherwise stated, all models include drive-parent sex and Cas9 promoter as fixed-effect covariates, with a four-component random-effects structure: publication/observation, species (independent and phylogenetically correlated), and construct identity (see Methods; model selection in Supplementary Data S5). Additional factors of interest are added individually to this baseline.

The I^2^ index indicates that over 90.5% of variability in drive inheritance cannot be attributed to sampling error alone (total *τ* ^2^ = 1.6588; *Q_E_* = 38882.3, *p* = < 0.0001). Species is a strong predictor of drive performance (*Q_M_ p* = < 0.0001; Fig 1b), with *Anopheles* mosquitoes showing the highest predicted inheritance rates (*An. gambiae*: 95.1%, 95% CI 91.6%–97.2%; *An. stephensi*: 93.2%, 95% CI 83.9%–97.3%) and substantially lower rates in other Diptera species (*Ae. aegypti*: 66.9%, 95% CI 46.5%–82.4%; *Dr. melanogaster*: 75.8%, 95% CI 58.7%–87.3%), though with wide and overlapping confidence intervals. Laboratory strain shows substantial within-species variation (Fig S8), but formal modelling of strain effects is infeasible due to near-complete confounding with publication and design factors (Fig S9). Although the species effect is large, design features correlated with species could contribute to the observed differences.

**Figure 1:**
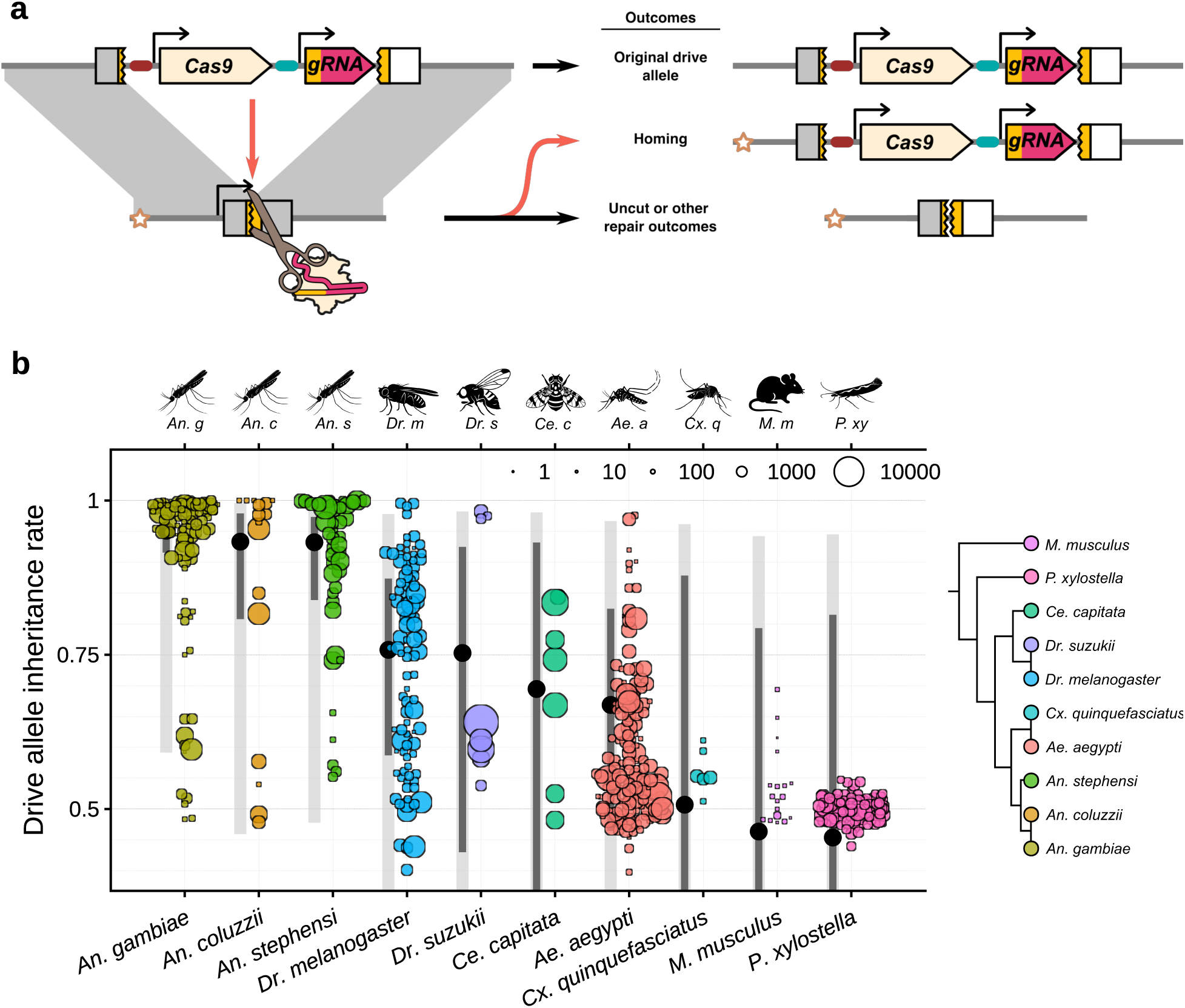
Homing gene drive schematic and per-species drive inheritance. **(a)** Overview showing a CRISPR-Cas9 homing gene drive converting a wild-type allele into a drive allele via inter-homologue gene conversion, alongside alternative outcomes (e.g., retention of the original allele or resistance allele formation). **(b)** Empirical drive-allele inheritance rates are shown for each species, with each point representing unique crosses (meta-data defined replicates merged). *An. coluzzii* is a member of the *An. gambiae* species complex. Point size scales with progeny scored. The colour legend indicates species phylogeny. Confidence intervals (dark grey lines) reflect precision of the predicted mean (black point); prediction intervals (light grey lines) reflect the expected range for a new observation.

A formal model-comparison hierarchy (Supplementary Data S5) decomposes this heterogeneity further. Adding species to the null model yields a more accurate model with ΔAIC = 21.3 (LRT *p* < 0.0001). Applying a phylogenetic framework further improves model fit (phylogenetic random effect preferred over IID by ΔAIC = 5.3), suggesting that species-level differences appear to be driven by ancestry. However, the estimated phylogenetic signal (100.0%) is at the boundary of the parameter space (the non-phylogenetic species variance converged to zero: 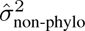 = 0.0000), and with only 10 species the exact partition of variance should be interpreted with caution. Nonetheless, this strong phylogenetic structuring of inheritance bias suggests that biological constraints dominate drive performance. To investigate the within species variability, we next evaluated specific design parameters, starting with the most frequently optimized factor: the timing of nuclease expression.

### 2.3 Nuclease expression timing is poorly predictive, and species-specific

The developmental timing of Cas9 expression, primarily controlled by the choice of promoter and 5*^′^*UTR, is widely considered a critical determinant of drive performance ^25;51^. Fig 2a shows drive inheritance rates stratified by the promoter element driving Cas9 transcription. We treated orthologous genes, whose promoter elements drive endonuclease expression, as equivalent across species (e.g., *mei-W68* and *spo11*), and found that adding promoter to a species-and-sex baseline reduces cross-to-cross variation by 10.2% (*τ* ^2^ reduction; LRT *p* = 0.0042; Supplementary Data S5). However, allowing promoter effects to differ between species substantially improves model fit compared to assuming a single cross-species ranking (ΔAIC = 45.6), indicating that promoter rankings are largely species-specific rather than transferable (LRT *p* = < 0.0001). The wide confidence intervals for individual promoter estimates (Fig 2a) reflect the realities of an emerging field: although the dataset is large relative to the gene drive literature, spreading observations across 30+ distinct regulatory elements, multiple species, and diverse construct designs leaves relatively few independent replicates per promoter–species combination. Prediction intervals, which additionally incorporate between-construct and between-study variance, span nearly the full range of possible inheritance rates for most promoters — indicating that the choice of promoter, by itself, is a poor predictor of outcomes when applied to new experimental crosses.

**Figure 2:**
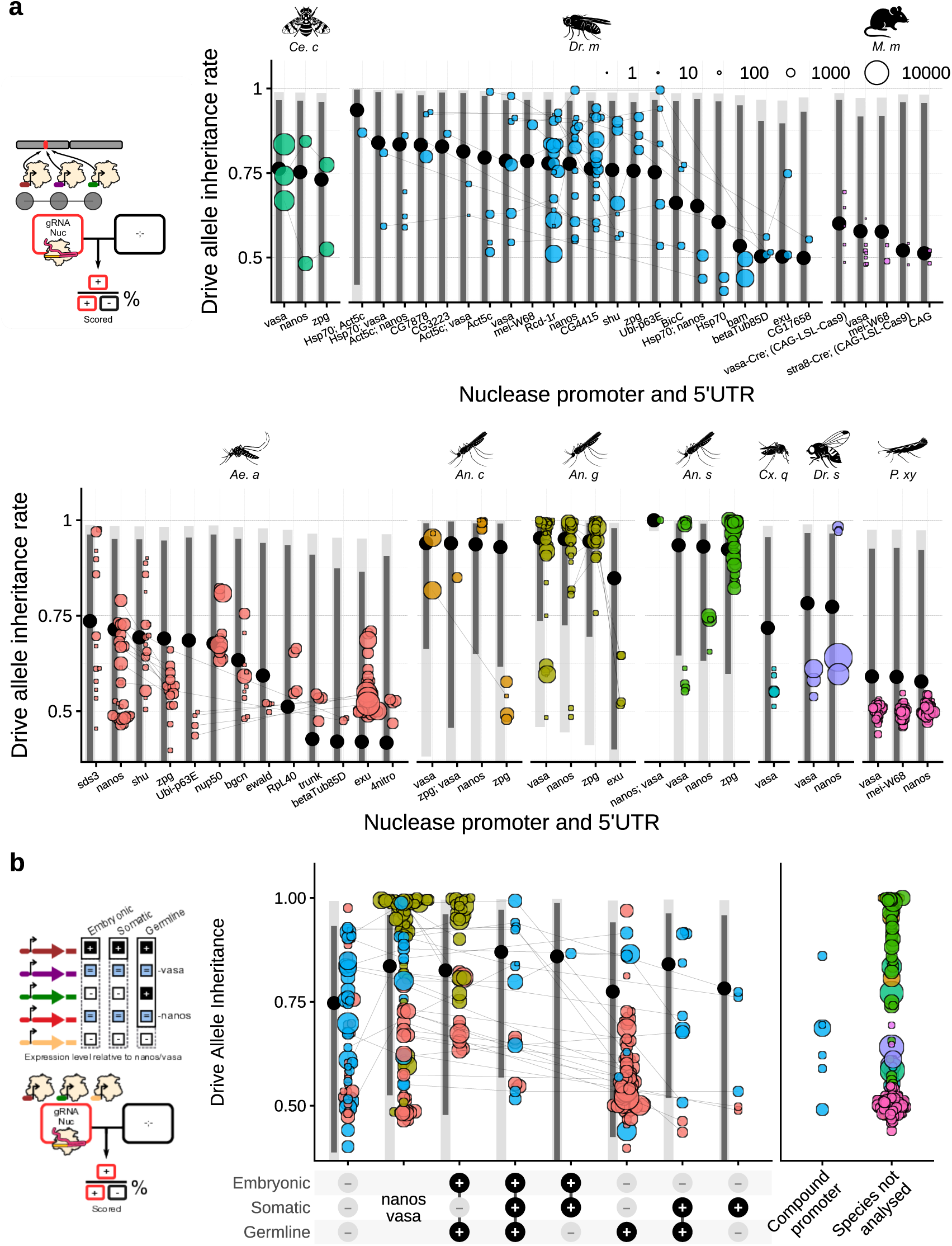
Promoter choice and expression-profile classification. **(a)** Drive-allele inheritance rates by nuclease regulatory module (promoter and 5*^′^*UTR) stratified by species. **(b)** Drive-allele inheritance rates by *Dr. melanogaster*, *Ae. aegypti*, and *An. gambiae* expression profiles of the endogenous gene of the nuclease promoter relative to nanos/vasa across developmental stages. Point size scales with progeny scored. Confidence intervals (dark grey lines) reflect precision of the predicted mean (black point); prediction intervals (light grey lines) reflect the expected range for a new observation. Grey line segments connect crosses equivalent apart from the Cas9 promoter used. ^11–21^

To move beyond simple promoter identity, we classified promoters by their gene-of-origin expression profiles using various bulk transcriptomics data spanning embryonic, juvenile, and adult stages, together with single-cell RNA-seq of the adult germline, for *Dr. melanogaster* (Fig S10), *Ae. aegypti* (Fig S11), and *An. gambiae* (Fig S12). Expression-profile class was a significant predictor (*Q_M_ p* = 0.1125; *n* = 425), but the two best-represented profiles — more germline-restricted than nanos/vasa (E−S−G+: 77.5%, 95% CI 42.5%–94.1%) and more broadly expressed (E+S+G+: 87.0%, 95% CI 56.8%–97.2%) — yielded comparable predicted inheritance rates.

We also investigated other nuclease expression and coding sequence factors. Neither promoter length (slope = 0.0000, *p* = 0.8591), 3*^′^*UTR length (slope = -0.0004, *p* = 0.0783), nor Cas9 codon optimisation (*Q_M_ p* = 0.4214; no significant differences between pairs) were significant predictors (Fig S13a-c). Cas9 variants featuring nuclear localization signals (NLS) at both the N and C termini (NLS-Cas9-NLS) showed higher rates of inheritance (70.7%, 95% CI 38.6%–90.2%) compared to Cas9-NLS (60.9%, 95% CI 26.4%–87.1%), but the difference was not significant (*p* = 0.5377) and is confounded with species (Fig S13c). The 3*^′^*UTR identity is heavily confounded with promoter and 5*^′^*UTR use (Fig S14), and cannot be independently assessed.

### 2.4 Substantial between-site heterogeneity implicates genomic context

While promoter choice is the most common purposefully varied parameter, specific combinations of transgene design choices may also influence drive performance. The core model includes a construct-identity random effect that captures substantial between-construct variance (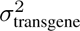 = 0.3518; boundary-corrected LRT *p* < 0.0001; ΔAIC = 104.0; Supplementary Data S5).

We attempted to decompose this construct-level variance into sub-components by testing whether broader groupings explain some of the construct-level variance, overlaying these over `transgene_ID` (Supplementary Data S5); the hierarchical nesting of design choices within publications is shown in Figs S15 and S16. Gene drive target gene (Fig S17) within each species did not account for a significant fraction of construct-level variance (0%; *p* 0.5000; ΔAIC = -2.0; 42% groups containing only one construct, mean 4.2 constructs per group). For transgene insertion locus (Fig S18), the test was likewise non-significant (1%; *p* 0.4797; ΔAIC = -2.0), but this result must be interpreted with caution: 57% of insertion-locus groups contain just a single construct (mean 2.0 constructs per group), leaving the test severely underpowered to detect between-locus effects. The between-construct heterogeneity thus cannot be attributed to any single coarser grouping tested here, though a contribution from insertion locus cannot be ruled out given the limited within-locus replication in the current literature.

Due to the frequent use of semi-random methods of transgenesis (e.g. transposon-based), the Cas9 insertion locus is not always known; by contrast mapping the driving element is straightforward because each corresponds to a unique gRNA site. The genomic distribution of loci for which homing was reported in our dataset is shown in Fig S17. Centromere proximity, which is known to correlate with meiotic crossover frequency ^52^, is not a significant predictor of drive element inheritance rates (Fig S19; slope -0.1282 logit units per relative unit, *p* = 0.7446).

### 2.5 The effect of the sex of the drive-carrying parent is complex and varies between species

Using within-study paired comparisons (Fig S9; see S2), we evaluated factors assessed for the same drive strains, beginning with the sex of the drive-carrying parent (Fig S20). Fig 3a contrasts inheritance from female versus male drive parents. Crosses performed in only one sex, commonly when the target gene is X-linked or females are sterile due to somatic cutting of specific target genes, appear along the axes but are not included in the statistical comparison. The overall difference was not statistically significant (OR = 1.23, 95% CI 0.91–1.66, *p* = 0.1715; *n* = 218). However, per-species tests reveal a significant female advantage in *Ae. aegypti* (OR = 1.21, *p* = 0.0274; *n* = 71) and a significant male advantage in *Dr. suzukii* (OR = 0.83, *p* = 0.0187; *n* = 3), albeit based on very few paired observations, while other species show no significant effect (Supplementary Data S4).

**Figure 3:**
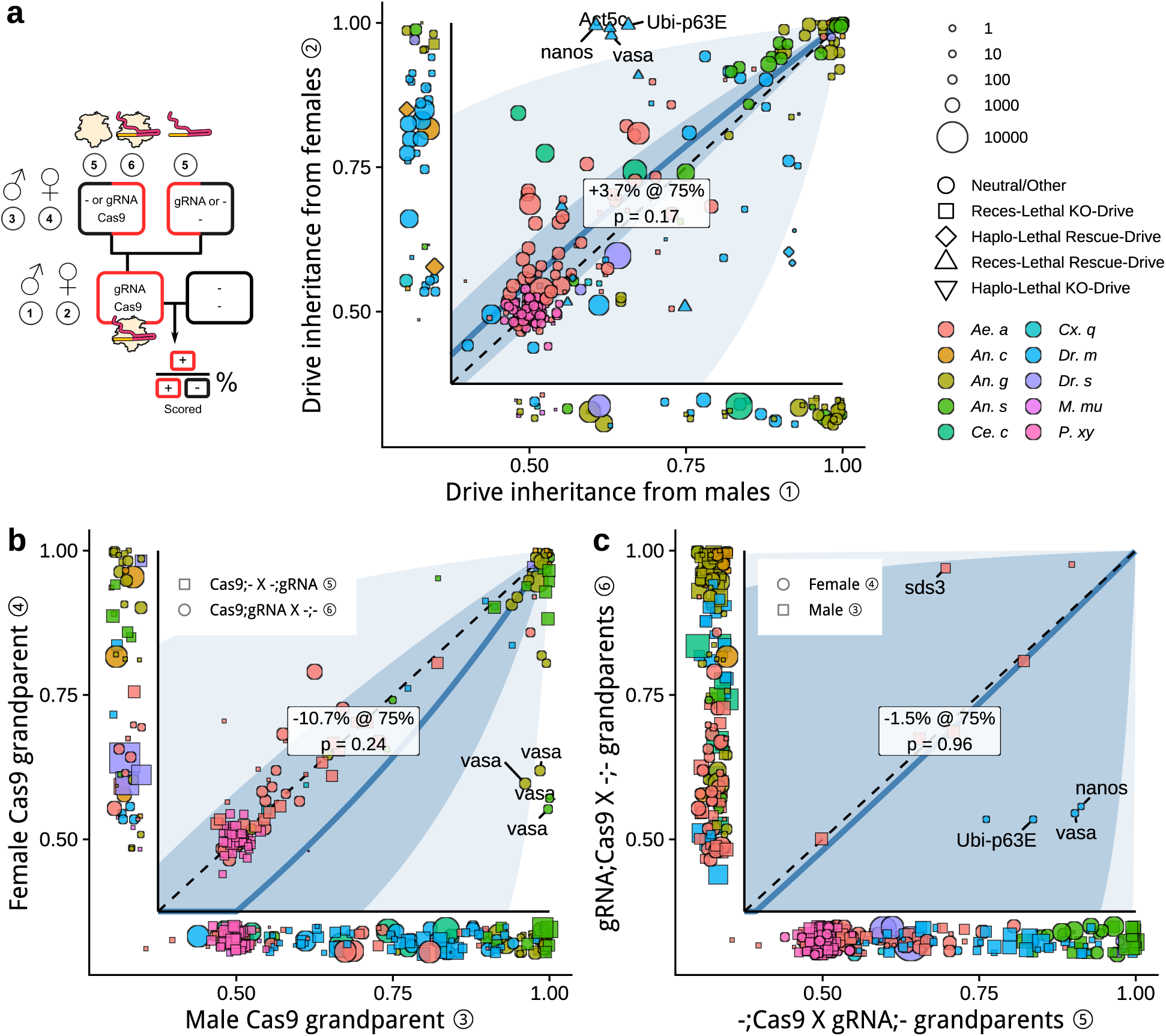
Drive inheritance across alternative parent-sex and deposition configurations. **(a)** Drive-allele inheritance from male (x-axis) versus female (y-axis) drive parents for paired crosses; point shape indicates target-gene fitness profile. **(b)** Drive inheritance when the Cas9-carrying grandparent was male (x-axis) versus female (y-axis); point shape indicates whether the Cas9 carrying grandparent also carried the gRNA. **(c)**: Drive inheritance when the Cas9-carrying parent lacked the gRNA (x-axis) versus when it also carried the gRNA (y-axis); point shape indicates grandparent sex. In all panels, unpaired crosses appear along the axes with jitter, and divergence from the diagonal indicates an effect of the contrasted factor. Point size indicates the lowest number of progeny scored in the two paired crosses. The fitted curve reflects the estimated log-odds ratio; inner ribbon = 95% CI (precision of the estimate), outer ribbon = 95% PI (expected range for a new paired observation). Dashed diagonal = no difference. Extreme values are labelled with the Cas9 promoter.

Significant outliers that imply higher rates of drive inheritance arising from female drive carriers were found to be associated with specific target-gene characteristics (Fig 3a, point shapes; Fig S21). This effect can likely be attributed to maternal deposition, where nuclease protein and/or gRNA expressed in the ovary can persist in eggs and affect the developing embryo in the following generation. In cases where the target gene is essential or haplolethal, drive inheritance can appear to be higher from females because maternal deposition and associated target-gene cleavage in the embryo culls non-drive offspring, whereas drive inheriting offspring survive through a drive-linked protective allele ^15^. However, the majority of drives in our dataset do not target essential genes, nor do they carry a rescue copy of their target gene.

We contrasted drive inheritance from individuals that themselves had a drive-carrying father or mother (grandparent generation). The overall grandparent-sex contrast was not significant (OR = 0.60, 95% CI 0.26–1.40, *p* = 0.2351; *n* = 121; Fig 3b, Fig S22). Per-species tests suggest an effect in *Anopheles*: in both *An. gambiae* (OR = 0.37, *p* = 0.0497) and *An. stephensi* (OR = 0.11, *p* = 0.0476), male grandparent Cas9 is associated with higher drive inheritance, with vasa-Cas9 as a pronounced outlier ^10^. Overall, data for *Ae. aegypti* revealed no significant effect of grandparent deposition (*p* = 0.5041; Supplementary Data S4). However, visual inspection suggests that maternal deposition may increase inheritance rates of poorly performing drives, while for performant drives maternal deposition is more likely to be detrimental.

We also evaluated the effect of the co-deposition of Cas9 and gRNA from the same grandparent compared to when they are contributed separately by one grandparent each (Fig 3c; Fig S23). We found no significant effect (*Q_M_ p* = 0.9583; *n* = 10), with only three studies directly contrasting these configurations (and in *Aedes* only evaluating paternal deposition) ^15;22;24^. In autonomous drives, Cas9 and gRNA are always inherited together, making it impossible to separate their contributions; in split drives, the extra generation needed to establish double-heterozygous grandparents limits the available data.

Biased inheritance caused by deposited endonuclease acting upon a drivable element in the absence of an inherited Cas9 transgene, a phenomenon sometimes referred to as “shadow drive” in the literature, can be assessed in non-autonomous drives that lack an expressed nuclease component (Fig S24). Shadow drive had not been detected in *Ae. aegypti* but had been observed in *Dr. melanogaster*, where it appears higher when Cas9 is deposited without the gRNA ^53^ and when the recipient is female ^15^. This suggests that the higher point estimate for females in Fig 3a may partly reflect intrinsic sex differences in DNA repair processes rather than transgene expression alone.

### 2.6 Somatic phenotypes reveal the reach of endonuclease activity beyond the germline

The modest effect of deposition on inheritance we detected may partly be an artefact of experimental design. Where loss of the target gene produces a visible phenotype, drive-carrying parents exposed to maternal deposition can be identified by somatic mosaicism (e.g., mosaic eye colour when targeting *white*). In the few crosses where distinct drive parent somatic phenotypes were reported, deposition-affected parents showed decreased drive performance in *Anopheles* but increased performance in *Ae. aegypti* (Fig S25). Crucially, researchers may select phenotypically uniform parents — either intentionally or because fitness costs eliminate heavily affected individuals — obscuring the true impact of deposition. We included drive parent somatic phenotype as a pairing criterion, leading to deposition effects only being contrasted between drive parents with the same reported somatic phenotype (see Discussion).

To further dissect the relationship between sex and nuclease activity, we analysed somatic phenotypes in F2 progeny (Fig S26; Fig S21), a smaller dataset that requires subdivision by additional experimental factors (see Methods; Fig S27). Within these groupings, somatic phenotype rates tend toward extremes — near 0% or near 100%. Important for the interpretation of the somatic phenotype data was recording F2 sex (Fig S28) and inheritance of the nuclease transgene, which in split drives segregates at Mendelian rates (Fig S29).

In contrast to the female bias in drive inheritance, we found no sex bias in the manifestation of somatic phenotypes among F2 progeny (Fig S30a; OR = 0.99, *p* = 0.9544; *n* = 111). However, somatic phenotype rates were markedly higher in Cas9;gRNA F2 progeny than in −;gRNA siblings (OR = 23.98, *p* = 0.0049; *n* = 108), indicating widespread somatic expression of Cas9 and demonstrating that many constructs do not have the desired germline-restricted autonomous expression, especially outside *Anopheles* (Fig S30b).

Somatic phenotype rates showed a large increase when the Cas9-carrying *parent* was female (OR = 22.91, *p* = 0.0675; *n* = 180), with the effect size far exceeding the corresponding maternal deposition effect on drive inheritance, although the overall test did not reach statistical significance due to high between-study variance. This asymmetry — large maternal effects on somatic cutting but modest effects on germline homing — suggests that deposited nuclease primarily affects the tissues in which it acts, but without altering the types of repair outcomes that determine inheritance in the poorly performing drives that dominate the dataset.

### 2.7 gRNA design features

We next evaluated gRNA-related design features. Multiplexing, i.e. the use of multiple gRNAs directed against the same target gene, has been proposed to improve drive robustness ^54^ but has limited representation in the dataset; the overall test was not significant (*Q_M_ p* = 0.2827), and the comparison is confounded with species and laboratory (Fig 4a). In contrast to the variety of endonuclease promoters used, the range of gRNA promoters used is narrow and confined to RNA polymerase III promoters derived from varying paralogs of the U6 and 7SK small nuclear RNA genes. The choice of gRNA promoter is heavily confounded with other lab-specific factors, and apparent performance differences should therefore be interpreted with caution (Fig 4b). Modified gRNA scaffolds have been adopted along-side multiplexing to reduce internal sequence homology and preclude undesirable recombination events in complex gene drive constructs. However, the co-occurrence of scaffold changes with multiplexing prevents us from evaluating the individual contribution of each in our dataset (Fig S31).

**Figure 4:**
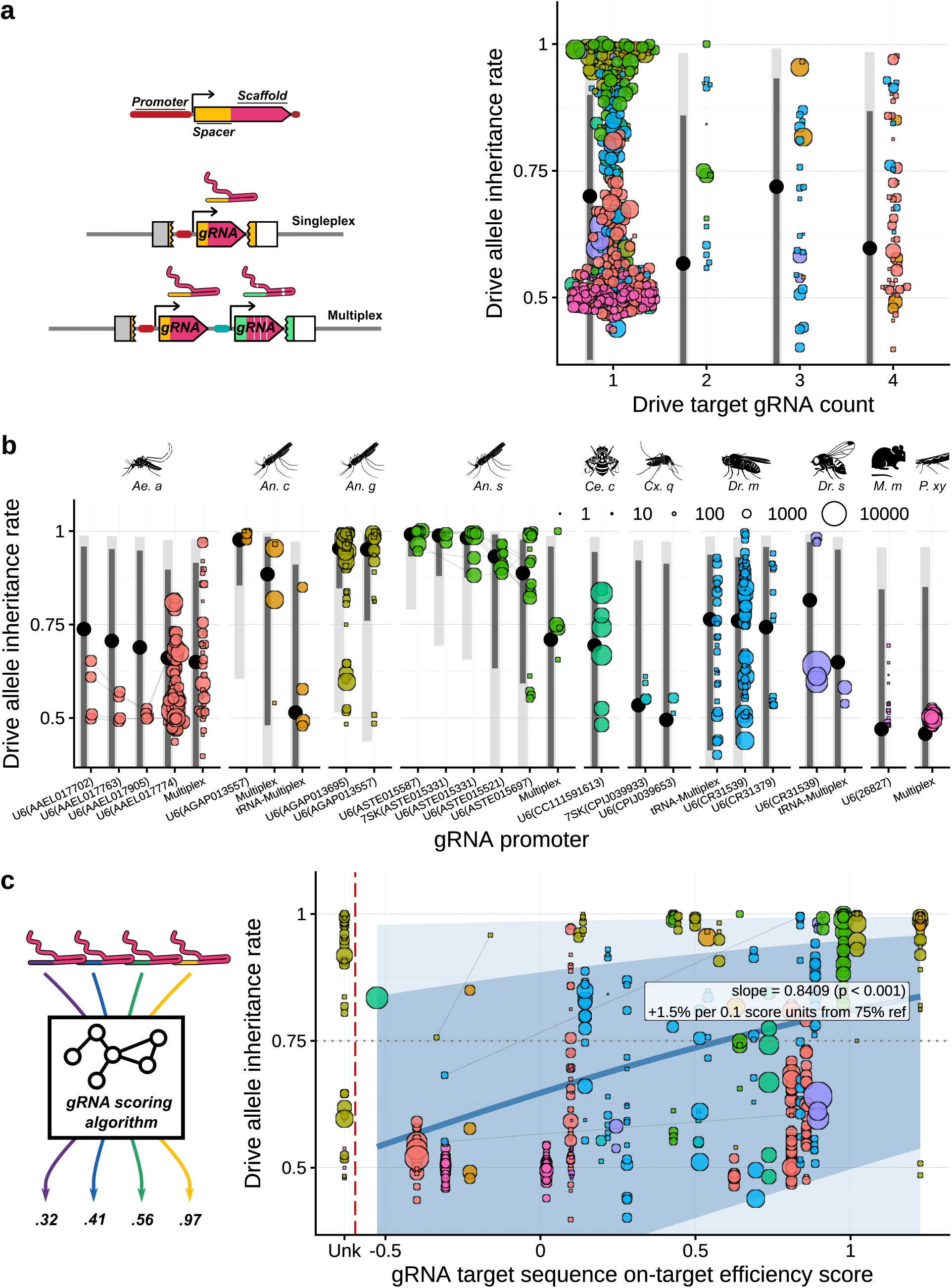
gRNA design features associated with inheritance bias. Inheritance rates are shown as a function of **(a)** the number of target-site gRNAs in the construct (multiplexing), **(b)** the regulatory element used to express the gRNA(s) ^24;43^, and **(c)** predicted on-target activity of the target-site gRNA (efficiency score) ^16;29^. Point size scales with progeny scored. Points are stratified by species. Grey line segments connect crosses equivalent apart fro_1_m_1_ the comparison factor. Cross-matching adjusted (visual impact only): gRNA target site, gRNA target sequence, target gene, and F2 phenotype gene excluded for the multiplex and gRNA promoter panels (confounded with comparison factor); gRNA target sequence excluded for the efficiency score panel.

On-target gRNA efficiency (RuleSet3 score) was a significant predictor (Fig 4c; slope = 0.8409 logit units per unit score, *p* = 0.0003). As a negative control, we evaluated the predicted activity of gRNAs in the region flanking each target site (Fig S32). The mean flanking-candidate score, reflecting local sequence context rather than selection, showed no significant association with inheritance (Fig S33).

### 2.8 Homing gene drive inheritance is robust in the face of DNA repair challenges

Finally, we examined whether physical features of the DNA repair substrates can predict drive performance. The effective resection distance, i.e. the length of recipient-chromosome DNA sequence that must be resected to expose homology to the drive allele, was not a significant factor (Fig 5a; slope = -0.0055 logit units per bp, *p* = 0.5806). Resection requirements above zero are concentrated in *Ae. aegypti* gene drive designs reaching a maximum required resection length of 63 bp. In the other species, well-aligned cut sites requiring little or no resection have been used predominantly, making this effect difficult to disentangle from the species effect. Nevertheless, a testable hypothesis is that reducing resection requirements could yield better performing gene drive designs in *Aedes*. For multiplexed designs, we assumed simultaneous cutting by all gRNAs (yielding zero resection when cuts flank the non-homologous segment); PAM-proximal and PAM-distal resection requirements appear comparable (Fig S34).

**Figure 5:**
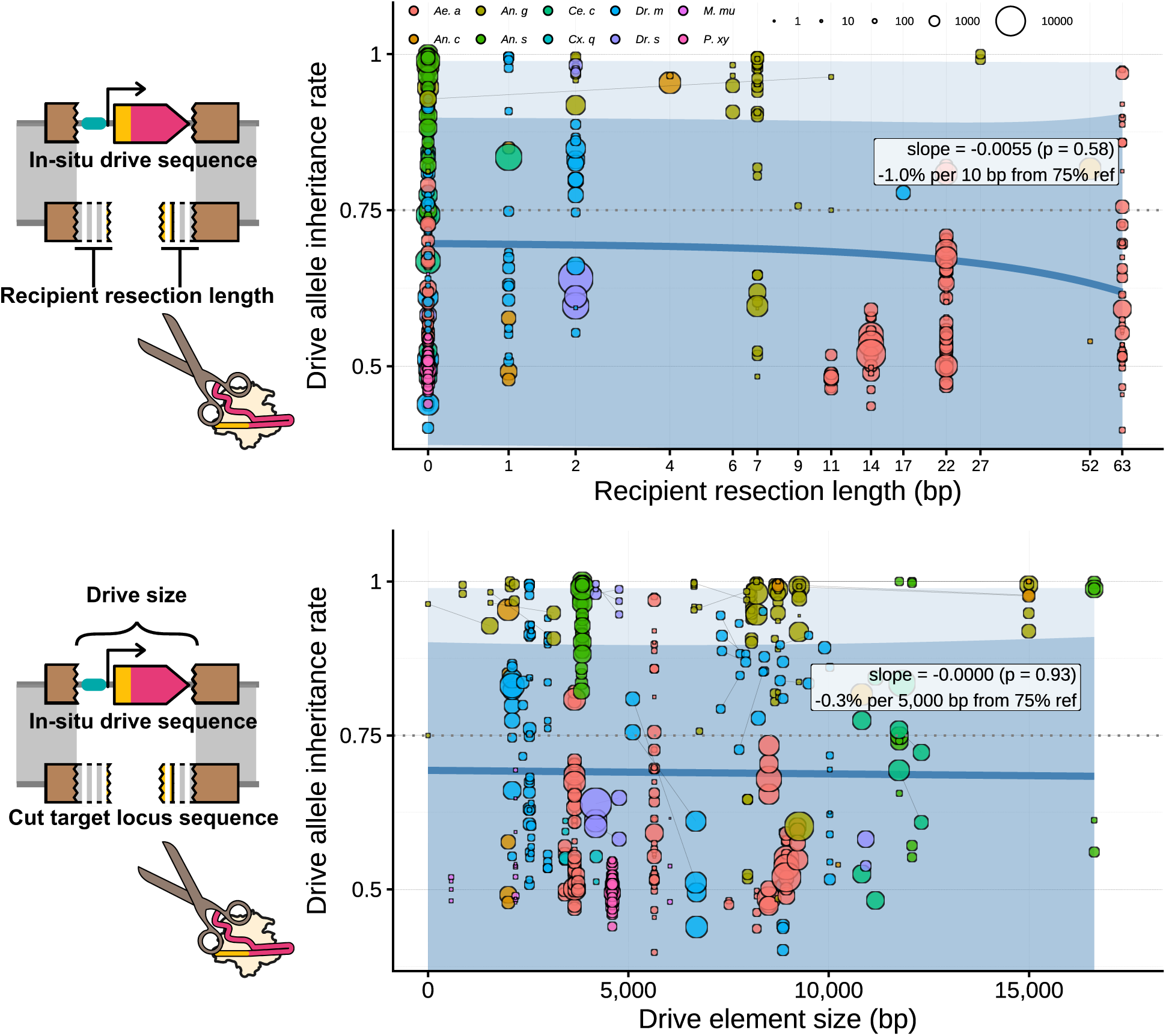
The effect of DNA-repair features on drive inheritance. Drive inheritance as a function of (top) the effective recipient resection length at the cut site ^36^ and (bottom) the size of the inserted drive element ^12;18;28;32;36;38;40;46^. Point size scales with progeny scored. Multiplex drives assume simultaneous cutting of all gRNAs (so could have 0 non-alignment). Grey line segments connect crosses equivalent apart from the comparison factor. Cross-matching adjusted (visual impact only): miscellaneous cross-level factor excluded for the drive-size panel because it often captures cargo presence, which co-varies with element size.

The drive alleles scored for inheritance bias in our dataset range from 0 bp in the form of a small deletion allele ^36^ (with Cas9 and gRNA expressed from separate elements) to 1°5 kb multi-effector autonomous drive elements ^9;28^. Surpris-ingly, drive element size was not predicted to affect gene drive performance (Fig 5b; slope = -0.0000 logit units per bp, *p* = 0.9332). The data include relatively well controlled paired-data in the form of conditionally removed marker genes and absence or presence of anti-pathogen cargo (Fig S35). This does introduce confounding by species, because anti-pathogen effector cargoes are commonly tested in *Anopheles*, which has a higher baseline rate of inheritance bias.

## 3 Discussion

Our cross-species meta-analysis identifies phylogeny as the strongest predictor of homing drive inheritance; separating the contribution of fundamental biology from the correlated design choices and genomic contexts in which drives have been tested remains a challenge. Between-construct variance is large, and attempts to attribute it to individual design factors — insertion locus or target gene — were inconclusive, suggesting that the full combination of design choices may determine construct-level performance. The overall effects of drive-parent sex and maternal nuclease deposition on inheritance are modest and species-dependent, whereas deposition has a pronounced effect on somatic phenotypes of the offspring.

The limited explanatory power of Cas9 promoter choice is perhaps the most practically relevant finding, given the field’s heavy investment in testing various germline-specific regulatory elements. Several factors may account for this. First, promoter identity is heavily confounded with species, laboratory, and other design choices. Second, the specific combination of transgene design parameters — including insertion site, target gene, and regulatory architecture — may collectively exert a stronger influence on HDR efficiency than nuclease expression timing alone. Third, our transcriptomic classification of promoters by expression profile similarly failed to explain substantial heterogeneity, suggesting that even biologically informed groupings do not capture the relevant axis of variation. These results do not imply that expression timing is irrelevant, but rather that its effect may be obscured by construct and locus-specific factors rarely varied systematically. Indeed, a finer-grained within-species analysis correlating single-cell transcriptomic profiles with the performance of 30+ Cas9 promoter constructs in *Dr. melanogaster* ^55^ found that optimal drive performance requires expression within a narrow cell-type-specific window, reinforcing that the relationship between expression level and drive outcome is not a simple more-is-better pattern and is unlikely to be captured by broad categorical groupings. Consistent with this, *in situ* hybridisation of Cas9 transcripts in *An. gambiae* gene-drive mosquitoes revealed cell-type-specific and sex-biased germline distributions ^56^, underscoring that spatial expression patterns not captured by bulk RNA-seq may be a functionally relevant axis. We hope our dataset can enable the development of more nuanced and predictive categorisations.

The expression classification itself carries methodological caveats that may contribute to its modest predictive power. Gene drive constructs use truncated regulatory fragments, not full endogenous promoters; missing enhancers or chromatin context may cause transgene expression to diverge from the native gene’s profile. Embryonic RNA-seq signal may reflect maternal mRNA deposition rather than zygotic transgene transcription, particularly for the *nanos*/*vasa* calibration genes which are known maternal contributors. Finally, the harmonization of expression data across diverse datasets introduces further biases. Expression categories should therefore be interpreted as approximate indicators of the promoter’s likely activity profile, not quantitative predictions of transgene expression.

More broadly, the pattern of multiple individually significant but weakly predictive factors reflects a structural feature of the gene drive literature. Species, promoter choice, insertion site, Cas9 coding-sequence variant, gRNA design and other aspects of construct design each capture partially overlapping sources of variation. Because most implementations require changing several parameters simultaneously, the variation explained by any one factor is rarely independent of the others. As we show, neither target gene nor insertion locus explains a significant fraction of between-construct heterogeneity, though limited within-locus replication leaves the latter test underpowered. The construct-level variance may be attributable to the full combination of design parameters, consistent with interacting contributions from genomic context, construct architecture, and species-specific biology. This shared variance structure means that ranking factors by individual effect size overstates the degree to which they represent independent optimisation axes.

Although the overall effect of drive-parent sex on inheritance was not found to be significant, per-species analyses reveal a significant female advantage in *Ae. aegypti*. The non-significance of the overall sex effect may partly reflect dilution by poorly performing drives, in which neither sex shows appreciable homing. Shadow-drive data in *Dr. melanogaster* indicating higher homing rates when the recipient of deposited Cas9 is female ^15;57^ point to an intrinsic sex difference in (germline) DNA repair outcomes rather than the effect of transgene expression alone. However, this sex-difference was not evident in our analysis of F2 somatic phenotype rates.

For maternal deposition, the per-species tests suggest an *Anopheles*-specific pattern: both *An. gambiae* and *An. stephensi* show higher inheritance when Cas9 is inherited from the male grandparent. *Ae. aegypti* shows no significant overall effect, which may be due to a beneficial effect of deposition at low baseline homing rates and a detrimental effect at high homing rates. The interpretation of these results depends heavily on whether the cut-rates of autonomously expressed Cas9;gRNA are limiting or not. Although it may seem that the cutting rate could readily be inferred from somatic phenotype rates in non-drive-carrying offspring, this proved more complex than expected. Interpreting somatic phenotypes in F2 offspring requires subdivision by a range of additional factors. The many required subdivisions fragment the data into small groups, within which phenotype rates tend toward extremes. More sophisticated models will be needed to separate the data into underlying parameters such as germline and somatic cutting rates and the contribution of maternal deposition.

An important caveat to our investigation of deposition is that pre-selection of parents with distinct somatic phenotypes could bias outcomes of the next experimental set of crosses. Researchers may, intentionally or through fitness-mediated selection, employ phenotypically uniform drive parent classes for their experiments. If deposition were to affect downstream inheritance differently in parents classed by their somatic phenotypes, preselection would obscure downstream effects on inheritance. We nonetheless included drive parent somatic phenotype as a pairing criterion, providing cleaner within-class comparisons at the cost of masking between-class deposition effects. We made this choice because in the few cases where drive parent somatic phenotypes are reported, they generally represent a deliberate investigation of this effect, and a disproportionate number of individuals with rare somatic presentations are scored, creating a biased view of deposition effects if they are lumped together. Put differently, a drive may look particularly susceptible to deposition only because researchers selectively investigated a minority of drive parents that presented with a strong somatic phenotype. Comparing only between parents with matched somatic phenotypes prevents this, at the cost of missing the impact of shifts in the ratio of parents with those distinct phenotypic classes because of deposition.

Our analysis has several limitations. First, we evaluated factors largely in isolation, but biological interactions are expected and correlation between research groups and design choices limits our ability to separate species-intrinsic biology from laboratory methodology; an expanded dataset, particularly from groups working across multiple species with shared designs, would help break these correlations. Second, our inclusion criteria are deliberately narrow: we required individual progeny counts from defined crosses, excluding cage trials, sequencing-only assays, and experiments where denominators could not be determined; parameters measured by other experimental designs (e.g., fitness costs from cage dynamics) should not be assumed absent simply because our analysis could not account for them. Third, drive allele inheritance rate is the net result of target-site cutting, HDR versus competing repair pathways, and differential offspring viability — not a single mechanistic parameter; indeed, distinct mechanisms can contribute concurrently to the observed bias ^58^. Our models treat the observed proportion as a composite outcome and cannot decompose it into constituent rates, though recombinatorily linked markers can in some cases distinguish homing from meiotic drive (Fig S36).

Our findings suggest where experimental effort in the gene drive field should be focussed. Construct-to-construct variation is substantial but could not be attributed to individual design factors using current datasets. Currently, insertion site is among the least systematically varied parameters in the literature — locus choice is typically constrained by other design goals (e.g. the repeated use of a small set of validated essential genes) — and could be an axis for further optimization of homing efficiency. The limited additional contribution of promoter choice, once correlated factors are accounted for, suggests that systematic experiments varying multiple design parameters together — including insertion loci and their interaction with regulatory architecture — deserve at least as much attention as promoter screening. Crucially, such multi-factor designs would provide the within-locus and within-target-gene replication needed to identify the sources of construct-level variation.

These recommendations are underscored by a further observation: even the near-perfect replication of experiments featuring the same drive construct can yield markedly different inheritance rates across studies. Our dataset contains nominally identical crosses from independent publications ^14;20;34^, i.e. crosses matched on all recorded metadata except the reporting publication (Fig S3). The outcomes in these identical crosses in *Anopheles* species span a range of inheritance rates far exceeding binomial sampling expectation. Similar within-construct variability is documented in *Aedes* ^24;25^, where replicate crosses of the same drive in different labs occurred through a different mechanism of inheritance bias. The replicate-cross analysis (Fig S7) confirms this more broadly: paired crosses that match on every recorded design parameter frequently fall outside the 95% binomial prediction intervals, indicating biological heterogeneity from factors not captured by any variable in our database. This irreducible variability sets a lower bound on the differences that any cross-study synthesis can reliably detect: if inheritance rates fluctuate substantially even when all *measured* parameters are held constant, it is unsurprising that comparisons across studies — where species, construct design, laboratory practice, and genetic background differ simultaneously — struggle to isolate the contribution of any single factor. We therefore arrive at the recommendation that future gene drive experiments report sufficient within-construct replication (multiple independent crosses per construct–sex–condition combination) to sufficiently characterise this intrinsic variability, and that study designs be powered to detect biologically meaningful differences above this background variation. Without such replication, the field risks continued investment in screening design variants whose apparent performance differences may be indistinguishable from the stochastic variation inherent to the system.

The structured dataset and analytical framework we provide are designed to support further investigation of these and other questions. For computational modellers, the machine-readable format enables direct parametrisation of gene drive models with empirical data, including parent-by-parent variability rarely incorporated in current models. For molecular biologists, the interactive web tool enables evaluation of design features through paired-cross comparisons with adjustable matching stringency. We have also recorded DNA sequences of drive elements and target sites to support downstream bioinformatic analyses. We invite the research community to use and extend this resource as the field moves from demonstrating that homing drives can work toward understanding, quantitatively, why they work better in some contexts than others.

## 4 Methods

### 4.1 Literature search and study selection

We searched PubMed and Web of Science (Supplementary Data S1), returning 3,400 unique records (Fig S1). After screening, 42 publications were selected for data extraction. We excluded cell-culture studies, singlecelled organ-isms, and *Dr. melanogaster* publications using only vasa- and/or nanos-driven Cas9. Non-CRISPR homing drives (principally I-*Sce*I) were not included because inheritance rates were not reported in a compatible format ^3–5^.

For each publication, we read the full text and, where available, supplemental data files. For each recorded value, we included a brief justification (quote, reference, or deduction). Data were excluded when only summary statistics were reported and individual progeny counts could not be determined.

We focused on data derived from the scoring of progeny from defined crosses, as this is the most commonly reported and standardised measure of drive performance. Cross data were included if they met all of the following criteria:

- Heterozygous locus: At least two alleles are described for the target locus, one susceptible to nuclease-induced DNA damage and one resistant.
- Defined transgene: DNA damage at the susceptible allele is expected to be generated by the product of an integrated transgene. This excludes exogenously supplied nuclease components (e.g., micro-injection).
- Heterozygous cross: A genetic cross is performed that may result in inheritance bias of a defined allele through homing.
- Defined genotypes: Individuals of each sex in the cross have a defined genotype for the target allele and nuclease transgenes. This generally excludes multi-generational cage trials, and crosses of mixed populations.
- Germline assessed: The outcome of the DNA repair process is evaluated by scoring progeny of the individual in whose germline it is expected to occur.
- Countable: Individual progeny counts are reported, and an inference can be made about the allele status. This excludes cases of “zygotic homing” evaluated by scoring somatic conversion in heterozygotes.

Where a cross meeting these criteria was identified, additional crosses with the same transgenes that did not strictly meet the criteria were generally included if they provided informative baselines or reference points.

### 4.2 Database structure

Each row of the database represents a grouping of F2 progeny from a published cross that cannot be further distinguished by any of the recorded metadata fields (Supplementary Data S2). Progeny from a single class may span multiple rows if they differ in, for example, the presence of a specific NHEJ mutation. Design-level factors (e.g., promoter, target gene) are shared across all rows from the same cross and often within a publication. In rare cases, we partitioned a single reported cross when additional recorded factors permitted meaningful subdivision (e.g., separating by D1 somatic phenotype).

We record genotypes for three generations: the scored F2 offspring; the F1 parents — the drive-carrying parent (D1) and the non-drive parent (T1) — in whose germline gene conversion is expected; and the F0 grandparents (D0.Male, D0.Female, T0.Male, T0.Female), whose nuclease deposition may influence outcomes. These generational labels are relative to the scored F2s; the same individual can be an F2 in one cross and an F1 in another. In crosses where homing may occur at multiple loci simultaniously, homing at each loci is scored as a seperate cross.

The primary outcome is the drive allele inheritance rate: the proportion of scored F2 progeny inheriting the drive allele. We recorded over ten secondary metrics, though data are generally very sparse for these (see interactive web tool).

### 4.3 Cross pairing criteria

Many crosses in our dataset were designed as within-study comparisons in which a single factor is varied while all others are held constant. Pairing is restricted to crosses within the same publication to reduce the risk of undisclosed confounders. These paired comparisons are largely free of the laboratory- and design-level confounds that complicate between-study inference, and are complementary to the multilevel models, which use the full dataset but rely on statistical adjustment for confounders.

The standard pairing factors are listed in Fig S9. For the nuclease-induced somatic phenotype metric, additional subdivision is required and these are listed in Fig S27. Furthermore, the drive carrying parent sex is replaced by the sex of the nuclease-carrying parent, and grandparental pairing factors are removed. Several pairing factors do not currently vary in the dataset (e.g., there are no cases where nuclease activity is anticipated in the non-drive parent), but are included in anticipation of future data where they may be relevant. Factors were excluded from pairing based on judgement of biological relevance. Additional miscellaneous cross-level differences not captured by the primary moderators are catalogued in Fig S35.

### 4.4 Meta-analytic models

All analyses were performed in R ^59^; software versions are listed in Supplementary Data S2. Multilevel meta-analytic models were fitted using the metafor package ^60^. The I^2^ index was calculated following Nakagawa and Santos ^61^ using the orchaRd package ^62^. Marginal means (predicted averages for each factor level) were obtained using the emmeans package ^63^, and pairwise differences between factor levels were assessed with Tukey adjustment for multiple comparisons. Throughout, confidence intervals (CIs) reflect precision of the estimated mean, while prediction intervals (PIs) additionally incorporate between-study variance and indicate the range expected for a new observation; both use the *t*-distribution with model residual degrees of freedom.

#### Drive inheritance models

For each cross, we calculated the log odds of drive allele inheritance (Fig S5) and the corresponding sampling variance using `escalc()` in metafor (measure = “PLO”), with a continuity correction of 0.5 added to zero cells only. The core model included D1 sex and Nuclease Promoter/5*^′^*UTR as fixed-effect moderators, with a four-component random-effects structure: ∼ 1 | Publication*/*obsID (between-publication clustering and observation-level overdispersion), ∼ 1 | Species (independent species intercept), ∼ 1 | Species_phylo (phylogenet-ically correlated species variance, using a correlation matrix from the Open Tree of Life), and ∼ 1 | transgene_ID (construct-level clustering, defined by an eight-column interaction of species, target gene, nuclease gene, promoter, regulatory factors, gRNA promoter, nuclease insertion variant, and gRNA target sequence). The random-effects structure was selected by an incremental model-comparison ladder (Supplementary Data S5). Additional moderators were added individually to this core model; model comparisons used maximum-likelihood (ML) estimation and were assessed by likelihood ratio tests (LRT) and AIC, while final variance-component estimates used restricted maximum likelihood (REML). Heterogeneity explained by a moderator is reported as the proportional reduction in observationlevel variance (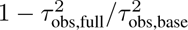).

Construct identity (transgene_ID) was modelled as a random effect — rather than a fixed effect — because the large number of unique constructs, many observed only once, precludes reliable fixed-effect estimation and the scientific question concerns generalisation to new designs; significance of variance components at the boundary was assessed by mixture *χ*^2^ LRT. The full model selection hierarchy, including sensitivity analyses for phylogenetic branch-length scalings and species subsets, is detailed in Supplementary Data S5.

#### Within-study paired comparisons

For factors varied within a study while all other design features are held constant (e.g., drive-parent sex), we calculated log odds ratios for each matched pair (measure = “OR”, with continuity correction added to all cells). These models used intercept-only fixed effects — confounders are controlled by the pairing design, and species heterogeneity is captured through the random-effects structure — with stepwise simplification of the random-effects structure in the rare cases when the full model fails to converge (Supplementary Data S2/S4). Per-species subgroup analyses were conducted where sample sizes permitted (Supplementary Data S4).

#### Somatic phenotype models

The number of F2 progeny displaying a knock-out or mosaic phenotype was used to calculate log odds and sampling variance. These analyses use F2-level data, requiring additional stratification by F2 sex, zygotic genotype, and nuclease transgene status beyond the cross-level variables used for drive inheritance. Progeny were grouped by cross and F2 zygotic genotype absent cutting, so that phenotype rates are calculated separately for F2 progeny with and without nuclease/gRNA transgenes.

### 4.5 Transcriptomic classification of Cas9 promoters

To evaluate whether biologically informed groupings of promoters explain drive performance, we classified the endogenous genes associated with each Cas9 promoter by their expression profiles. We obtained bulk RNA-seq data covering embryonic, juvenile, and adult life stages ^64–67^, together with single-cell RNA-seq of the adult germline ^68–70^, for three species well represented in our dataset: *Dr. melanogaster* (Fig S10), *Ae. aegypti* (Fig S11), and *An. gambiae* (Fig S12). Expression values were log-transformed and min-max scaled to [0, 1] independently per species and platform, then averaged across three developmental periods (Embryonic, Somatic, Germline). For simplicity, we use the term germline, but this includes some somatic tissues. Each gene–sex combination was classified as higher (+) or lower (−) expression levels in each period using thresholds calibrated from the reference genes *nanos* and *vasa*: the Germline threshold was set at the minimum of the two markers’ mean expression, and the Embryonic and Somatic thresholds above the maximum, so that the calibration genes themselves would classify as germline-restricted (E−S−G+). The resulting E±S±G± categories were used as a moderator in the meta-analytic model.

### 4.6 gRNA efficiency scoring

Predicted on-target gRNA efficiency was calculated using RuleSet3^71^. For multiplexed drives, we used the mean score across all target-site gRNAs.

### 4.7 Data validation

On multiple occasions, we contacted original authors for clarification when data were missing or ambiguous. Given the scale of the dataset, detailed external review may still identify issues in transcription, processing, or interpretation. We welcome these corrections and will maintain a post-publication public change log for updates.

### 4.8 Data and code availability

All supplemental data files (S1-S5), raw data, processed data, analysis scripts, and the interactive web tool are available at: https://osf.io/xcbpm.

### 4.9 Use of AI

Large language models were used during the preparation of this manuscript. Specifically, Anthropic’s Claude (Claude Opus 4.5/4.6 and related Sonnet and Haiku models) and OpenAI’s ChatGPT (GPT-5) were used to assist with writing and editing code, drafting and revising manuscript text. Species icons in figures were generated using OpenAI’s DALL-E. The authors reviewed and edited all AI-assisted output, and take full responsibility for the content of this publication.

## Acknowledgements

The authors thank Alessa Weiler, Joshua Ang, Tim-Harvey Samuel, Michelle Anderson and Luke Alphey. All authors contacted for clarification or additional data responded positively; we thank them for this. N.W. and S.A.N.V. were funded by the Gates Foundation grant INV-058071. N.W. and P.T.L. were funded by BBSRC grant UKRI1928. M.B.B. and S.A.N.V. were funded by BBSRC grants BB/H01814X/1, BB/L00948X/1, BB/V008110/1, BB/M011224/1.

## Author contributions

S.A.N.V. and M.B.B. conceptualised the study. S.A.N.V. led all investigations, data extraction, formal analysis, software development, visualisation, and writing. B.L. contributed to the bioinformatic analysis of drive and resection length. M.B.B., P.T.L., and N.W. provided supervision. P.T.L. and E.R.I. provided guidance on the statistical analysis approach. All authors provided comments on the manuscript draft.

## 5 Supplement

**Figure S1:**
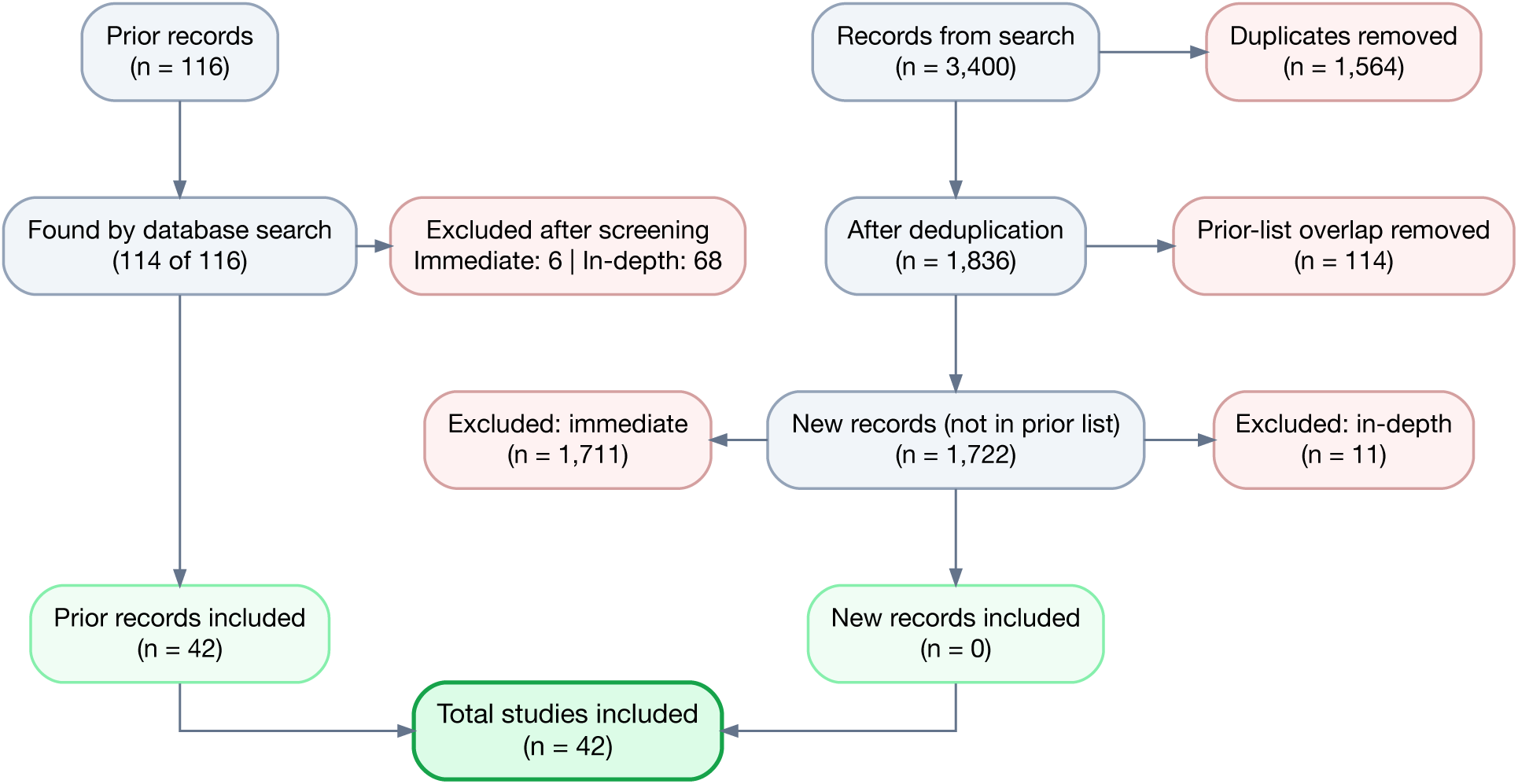
Literature search and screening flow diagram. PRISMA-style summary of prior records and database search yields, duplicate removal, exclusion categories, and the resulting set of records retained for full evaluation and data extraction. See S1 for more details.

**Figure S2:**
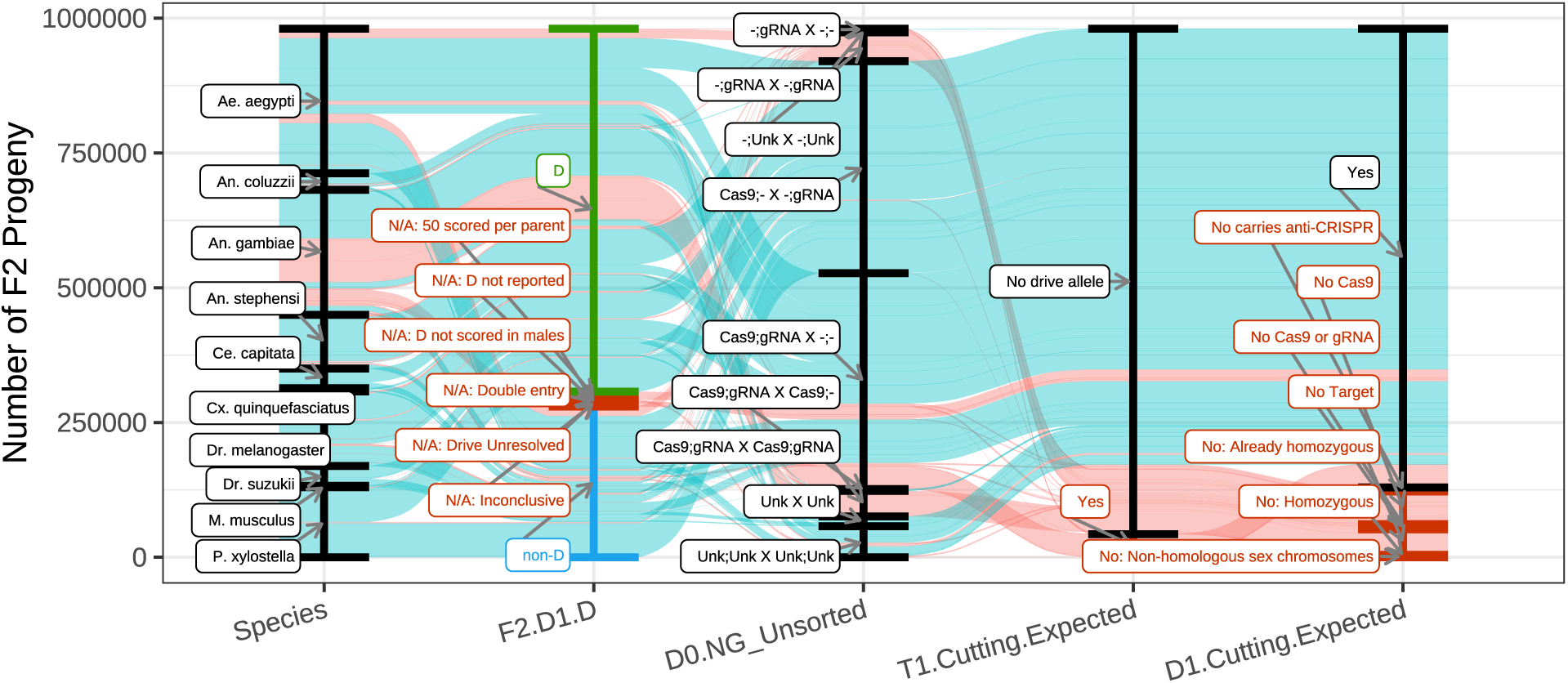
Filtering of progeny-counts records for the drive-inheritance metric. Sankey-style accounting of F2 progeny counts across sequential inclusion/exclusion criteria. Stream widths are proportional to progeny numbers, visualising where data are retained versus removed.

**Figure S3:**
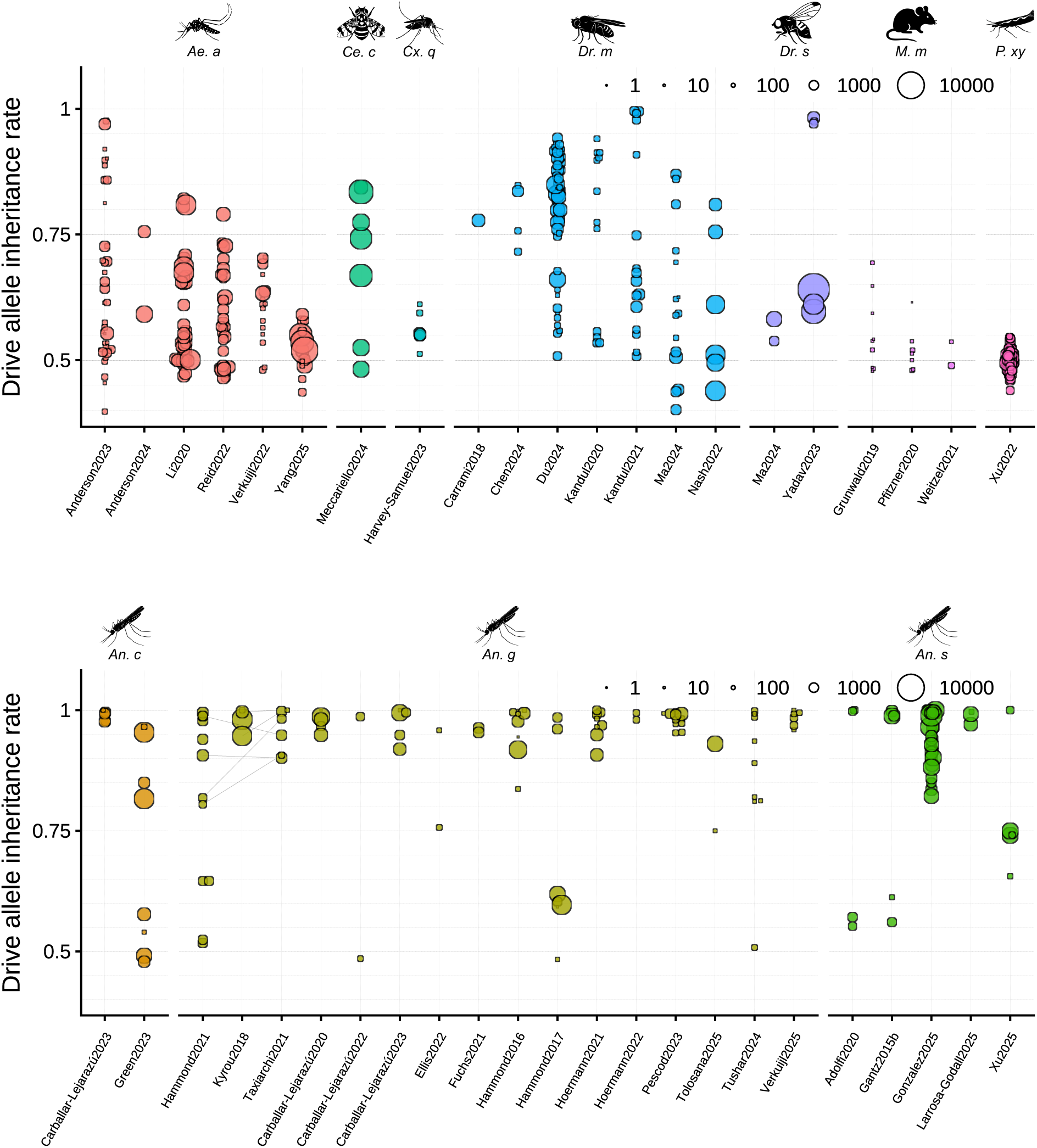
Drive allele inheritance rates by publication across species. The y-axis shows the drive allele inheritance rate, and the x-axis lists the source publication within each species panel. Point size scales with progeny scored. Crosses deemed equivalent within a publication by the metadata are merged; Grey line segments connect crosses equivalent apart from the publication of origin. ^14;20;34^

**Figure S4:**
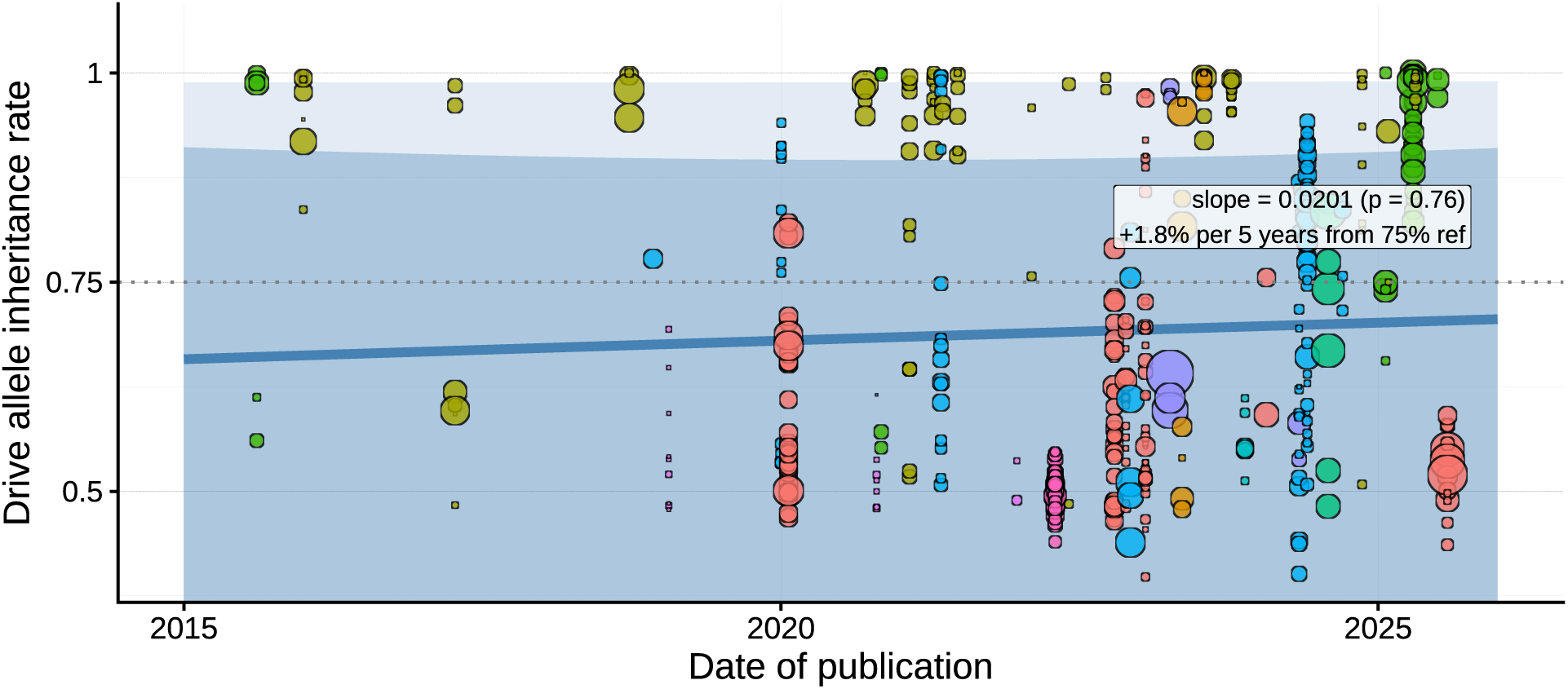
Temporal growth of the dataset. Timeline of included publications. The meta-regression line tests whether reported inheritance rates have changed over time.

**Figure S5:**
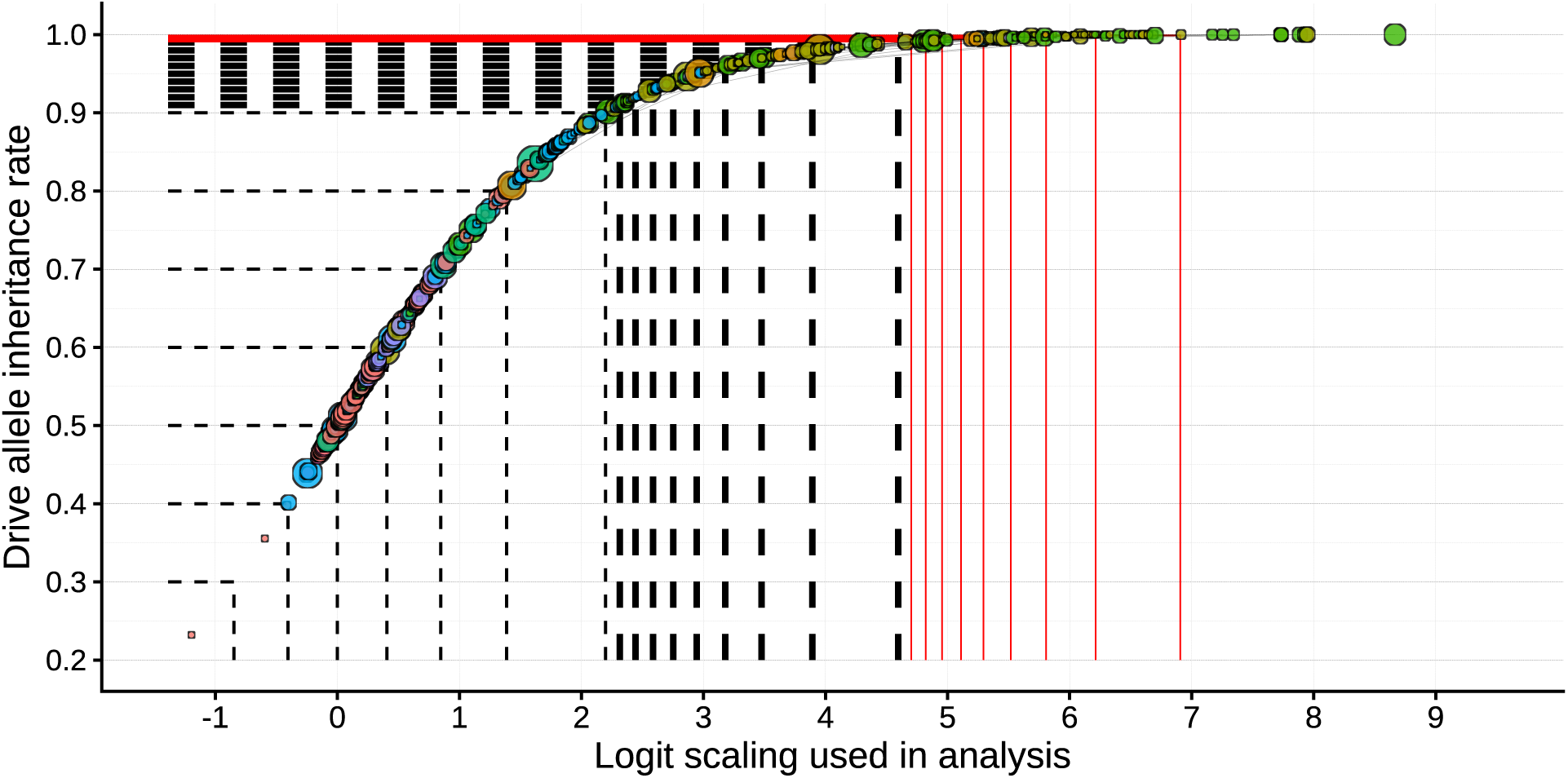
Logit transformation of inheritance rates. Mapping of raw inheritance fractions onto the logit (log-odds) scale used for meta-analytic modelling, illustrating the non-linear relationship near 1. Our statistical analysis assumes that the same effect size corresponds to a smaller change in inheritance % at extreme frequencies than at intermediate ones.

**Figure S6:**
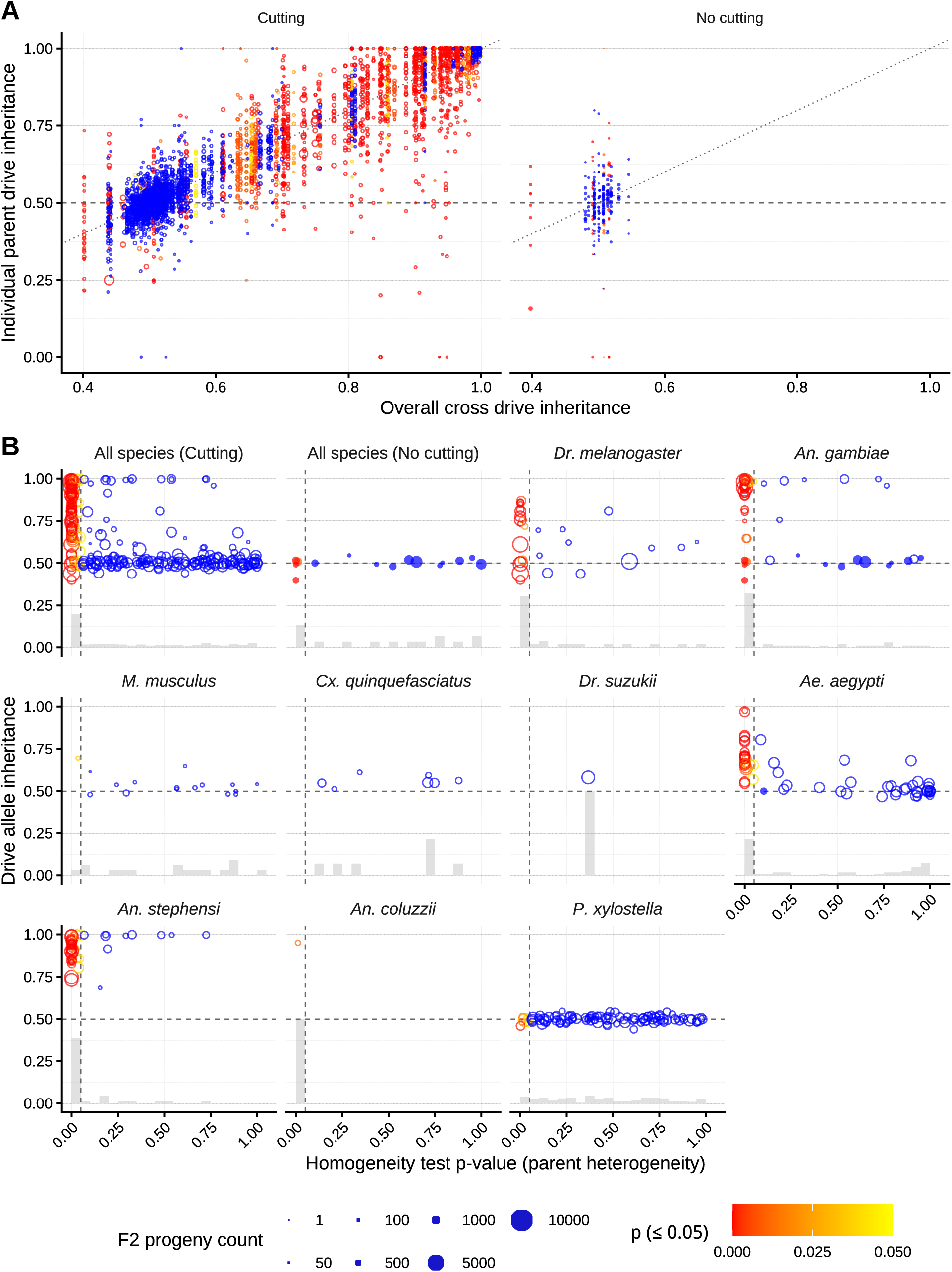
Excess parent-to-parent variability in inheritance rates. **(a)** For crosses with individual drive-parent data, each parent’s inheritance rate (y-axis) is plotted at the pooled cross mean (x-axis), stratified into cutting and no-cutting (control) crosses. In the cutting group, parent-to-parent values show extra-binomial variability (colour scale) when inheritance bias occurs. **(b)** For each pooled cross, a homogeneity-of-proportions test p-value (x-axis/colour scale) is plotted against overall inheritance (y-axis), faceted by species. Point size scales with progeny scored; colour indicates statistical significance (p *<* 0.05). The enrichment of low p-values in the cutting class supports that parent-to-parent variability exceeds binomial expectation specifically when drive activity occurs.

**Figure S7:**
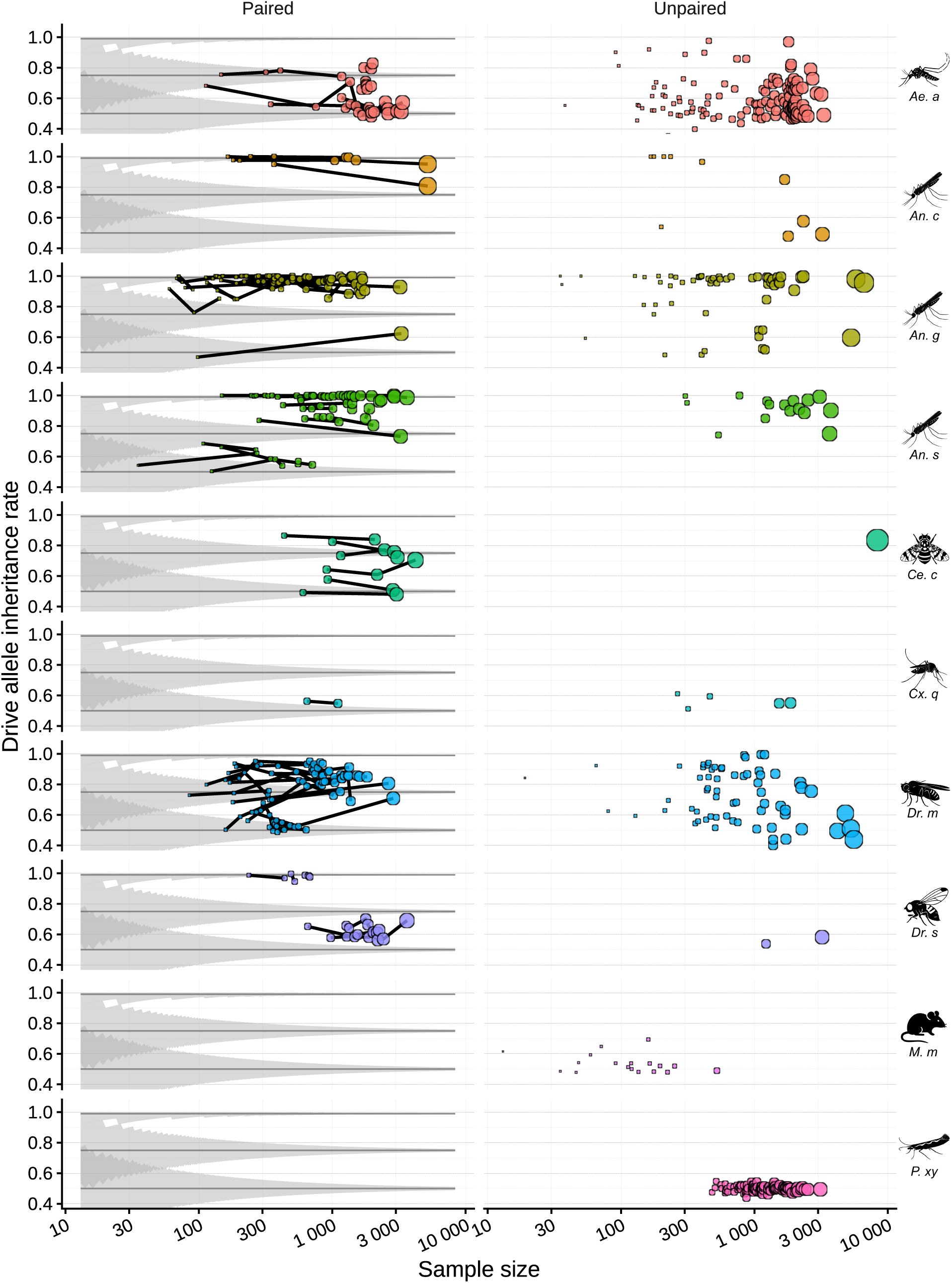
Consistency of drive inheritance across metadata-defined replicate crosses. **(a)** Crosses considered equivalent by the study metadata (Fig S9) are connected by line segments; in all other figures, these replicates are merged for visual clarity. Shaded funnels show binomial 95% quantile bands at true fractions of *p* = 0.50, 0.75, and 0.99. **(b)** Crosses with no equivalent replicate appear as unlinked points. Point size scales with progeny scored as indicated by the x-axis.

**Figure S8:**
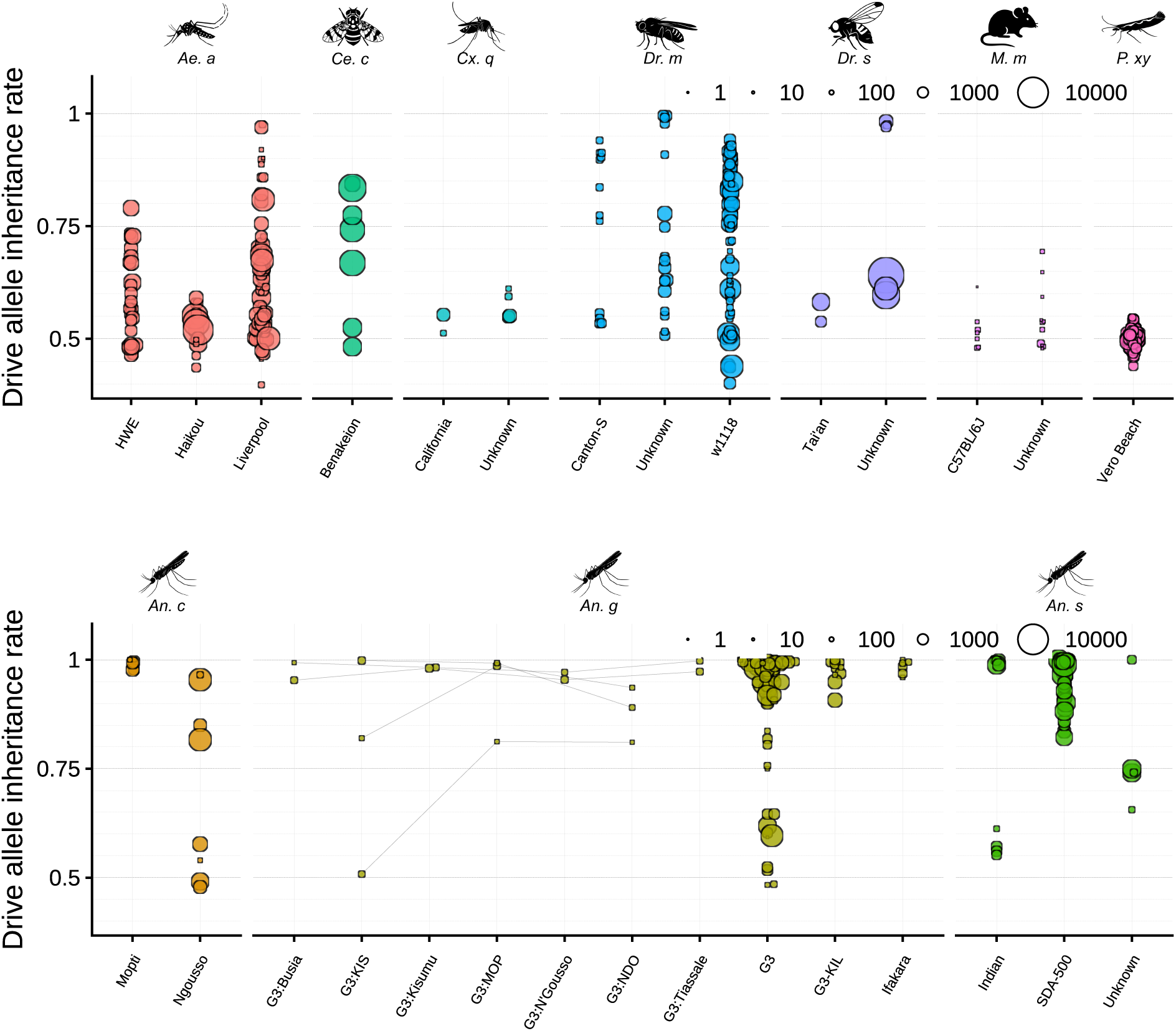
Drive allele inheritance by species strain. Observed drive allele inheritance rates are shown for individual experimental crosses stratified by reported strain and faceted by species. Point size scales with progeny scored. Grey line segments connect crosses equivalent apart from the laboratory strain. ^35;37^This is likely an underestimate of strain diversity, especially when not all strains are made *de novo* in the particular publication (e.g., imported from a strain repository). *An. coluzzii* is a member of the *An. gambiae* species complex, and G3 is a hybrid of the two.

**Figure S9:**
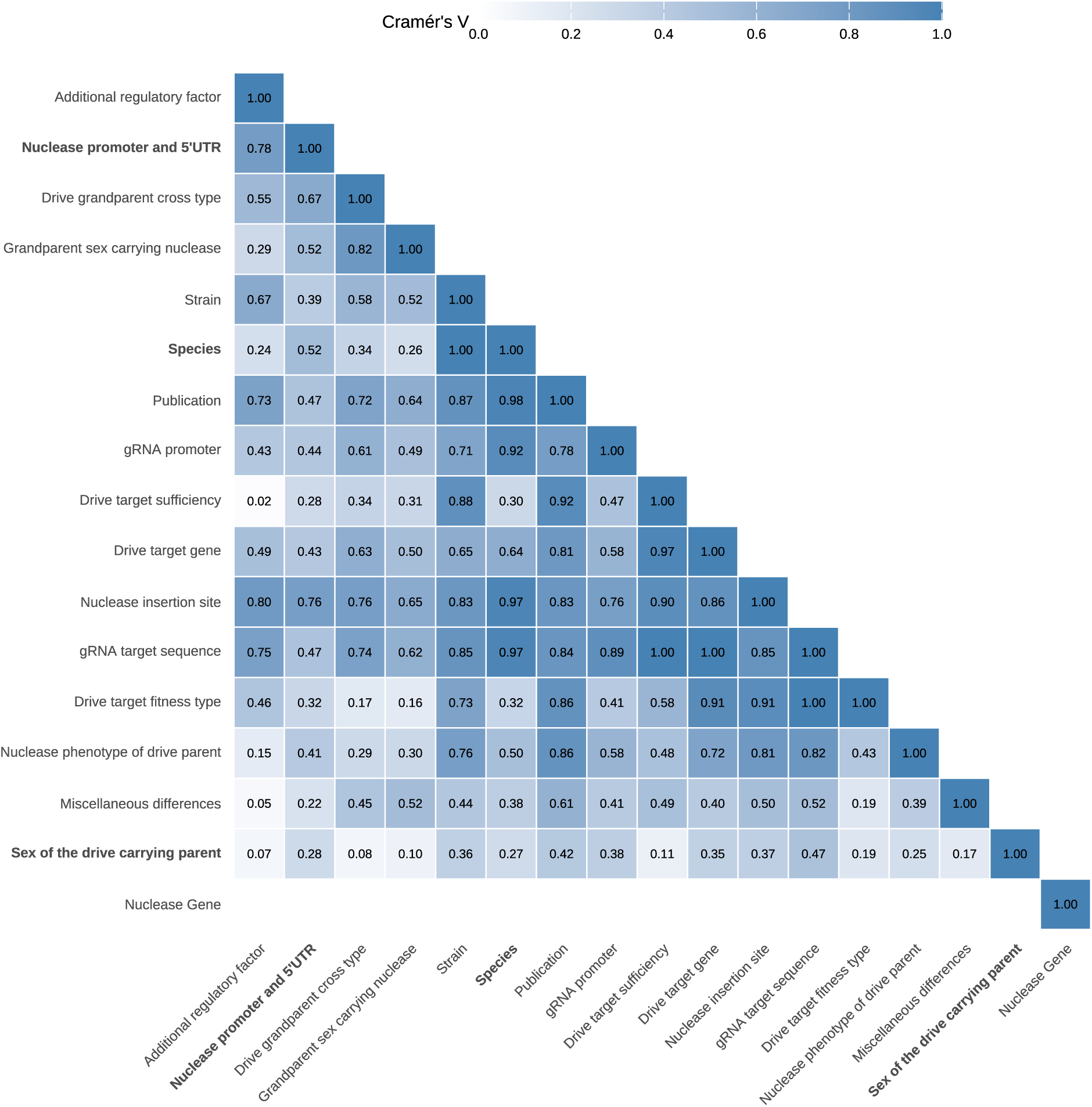
Correlation matrix of F1-level pairing factors for drive allele inheritance. Each cell indicates whether pairs of factors co-vary across the dataset. Factors that are highly correlated are effectively redundant for pairing purposes; factors that vary independently provide distinct axes of experimental control.

**Figure S10:**
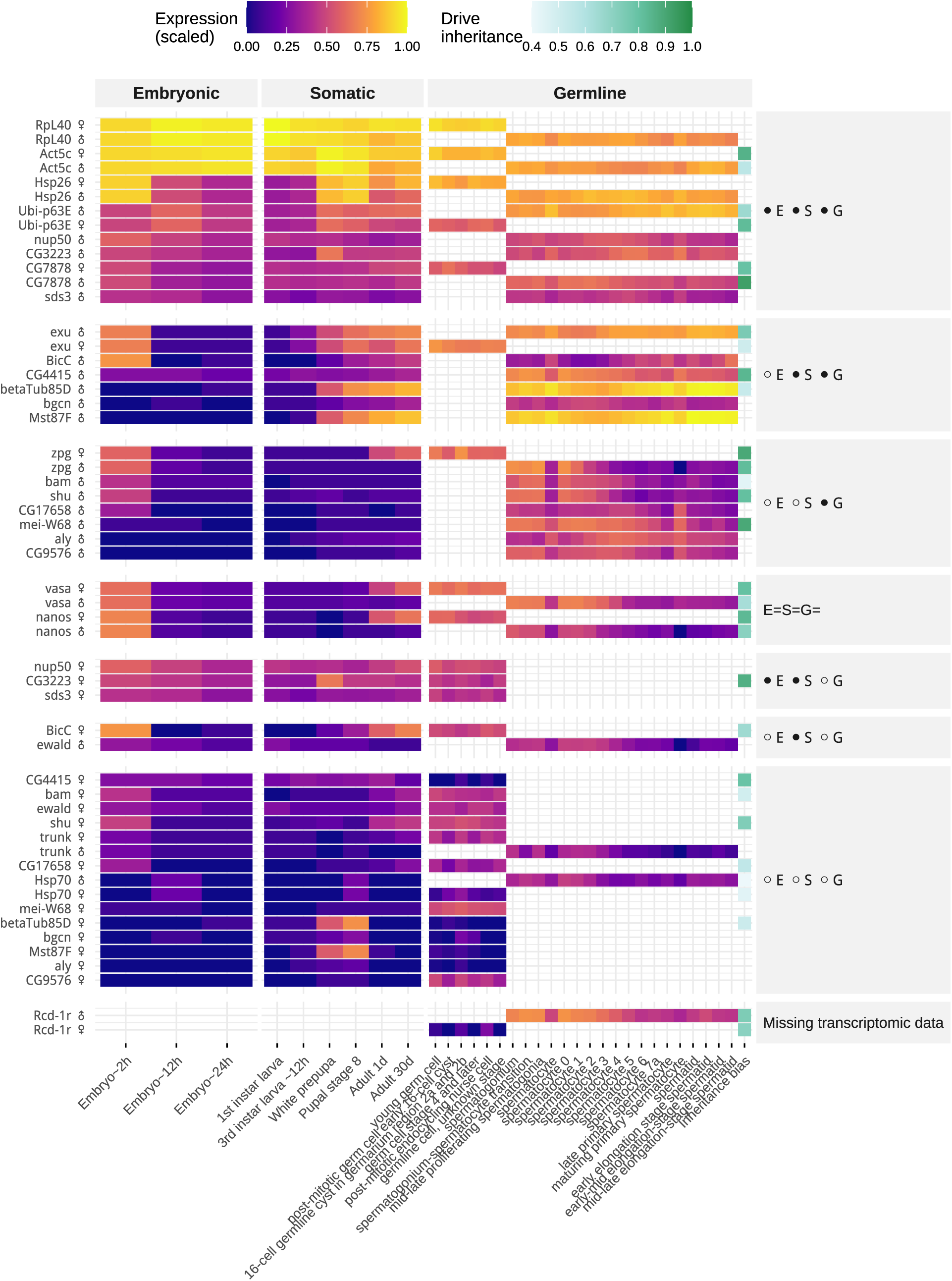
*Drosophila melanogaster* developmental and germline expression of genes associated with Cas9 promoters, alongside observed inheritance bias. Heatmap rows list promoter-associated genes used to drive Cas9 expression in homing gene-drive constructs (y-axis). Rows are stratified by sex. Columns summarise normalized gene-expression levels across annotated developmental timepoints (embryo to adult; left) and germline cell types. The far-right column reports the corresponding drive inheritance rate. White cells indicate missing/unavailable expression measurements for a given gene–stage combination.

**Figure S11:**
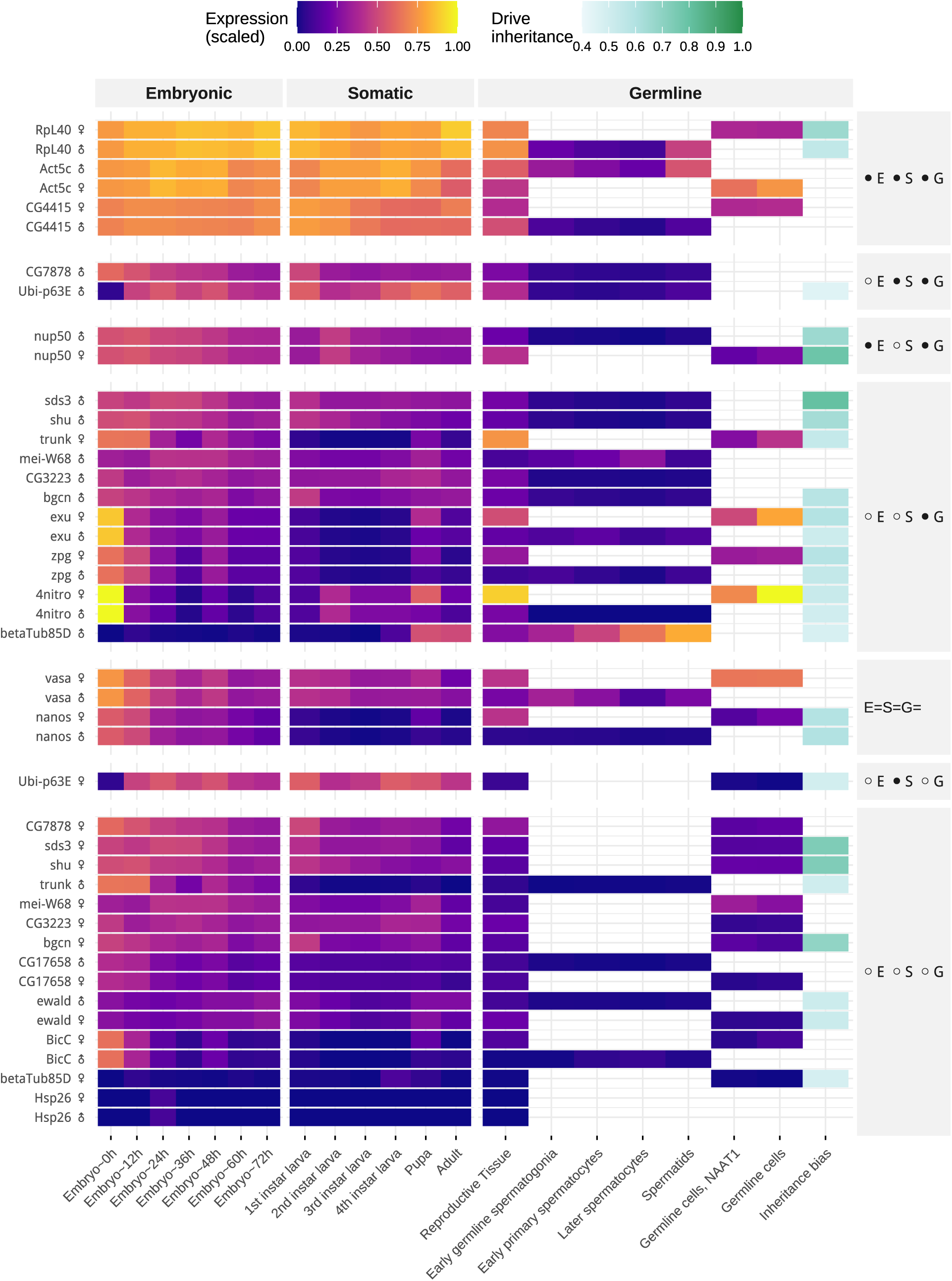
*Aedes aegypti* developmental and germline expression of genes associated with Cas9 promoters, alongside observed inheritance bias. Heatmap rows list promoter-associated genes used to drive Cas9 expression in homing gene-drive constructs (y-axis). Rows are stratified by sex. Columns summarise normalized gene-expression levels across annotated developmental timepoints (embryo to adult; left) and germline cell types. The far-right column reports the corresponding drive inheritance rate. White cells indicate missing/unavailable expression measurements for a given gene–stage combination.

**Figure S12:**
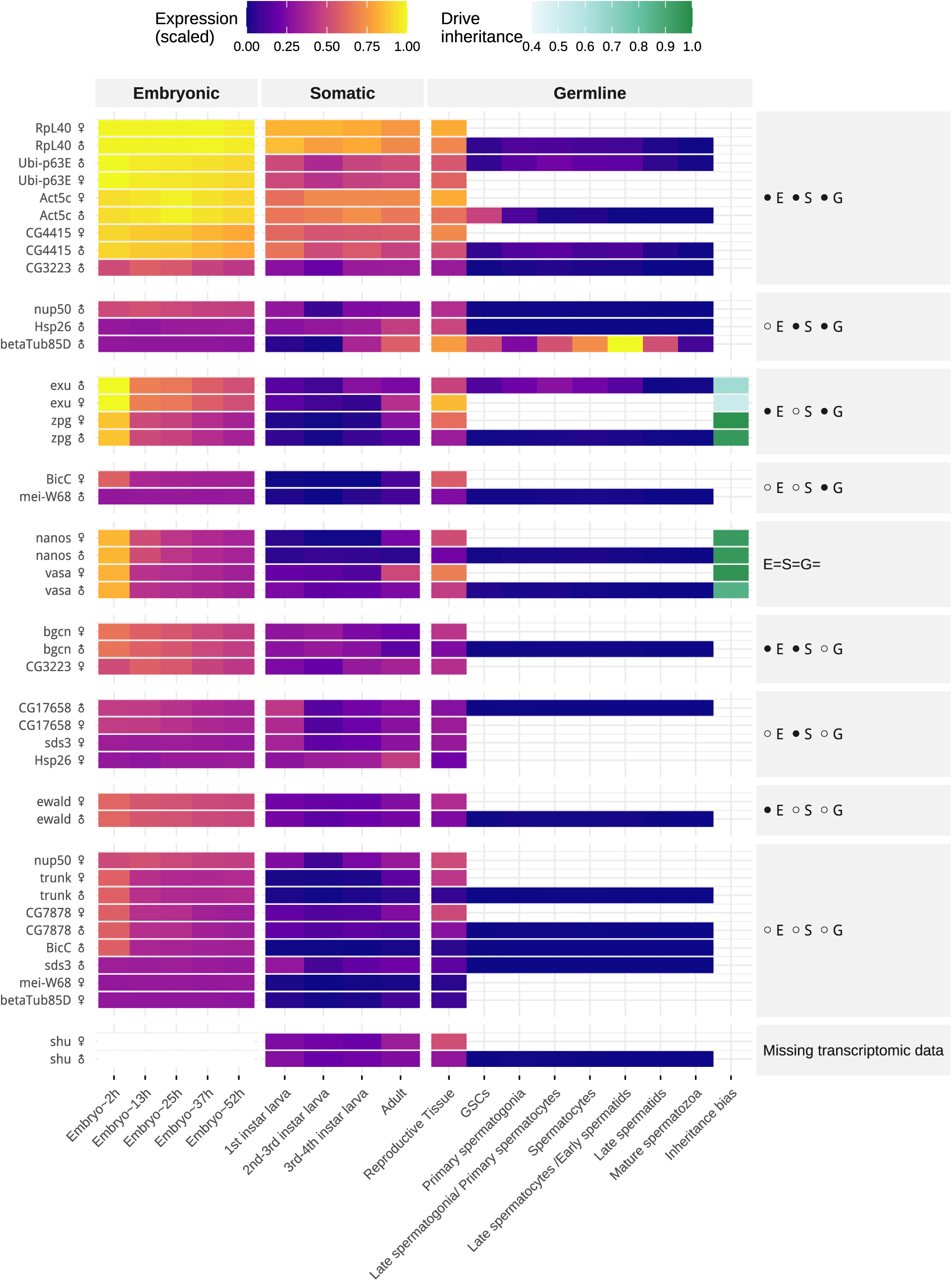
*Anopheles gambiae* developmental and germline expression of genes associated with Cas9 promoters, alongside observed inheritance bias. Heatmap rows list promoter-associated genes used to drive Cas9 expression in homing gene-drive constructs (y-axis). Rows are stratified by sex. Columns summarise normalized gene-expression levels across annotated developmental timepoints (embryo to adult; left) and germline cell types. The far-right column reports the corresponding drive inheritance rate. White cells indicate missing/unavailable expression measurements for a given gene–stage combination.

**Figure S13:**
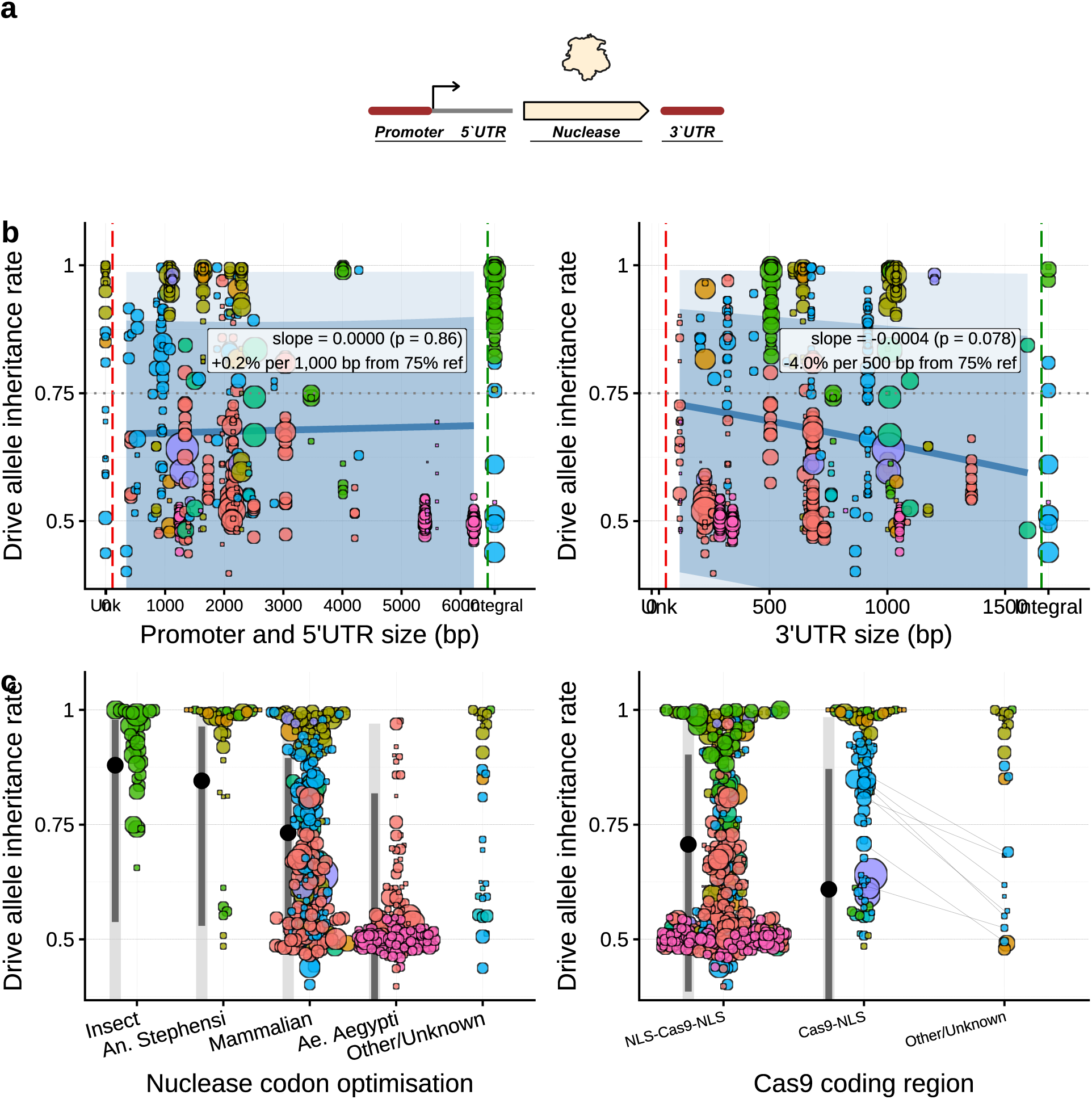
Drive inheritance with various Cas9 regulatory element features. **(a)** Schematic of the nuclease open reading frame, highlighting the regulatory and structural elements evaluated in subsequent panels. **(b)** Promoter length (left) and 3*^′^*UTR length (right) versus drive inheritance. Integral indicates elements that directly make use of the full endogenous gene’s regulatory context at the site of insertion. **(c)** Drive inheritance stratified by how the Cas9 coding sequence was codon-optimised (left) and by the number of nuclear localisation sequences (NLS) on Cas9^12^(right). Point size scales with progeny scored. Species are indicated by colour. Grey line segments connect crosses equivalent apart from the comparison factor.

**Figure S14:**
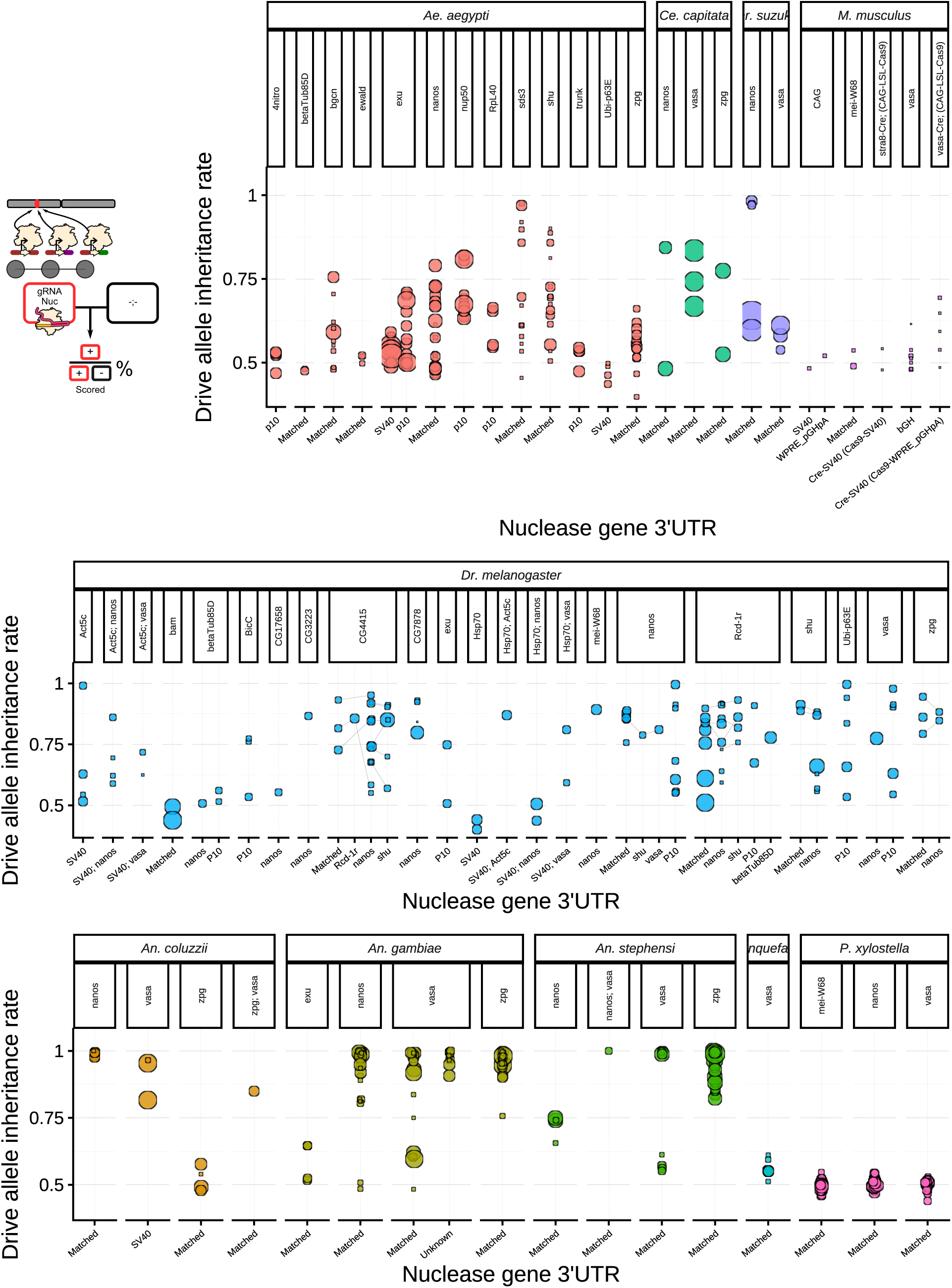
Drive allele inheritance rates by Cas9 3’UTR. Observed drive allele inheritance rates are shown, faceted by Cas9 promoter. Point size scales with progeny scored. Grey line segments connect crosses equivalent apart from the 3’UTR used. ^11;12^ ‘Matched’ indicates the 3’UTR is from the same gene as the promoter.

**Figure S15:**
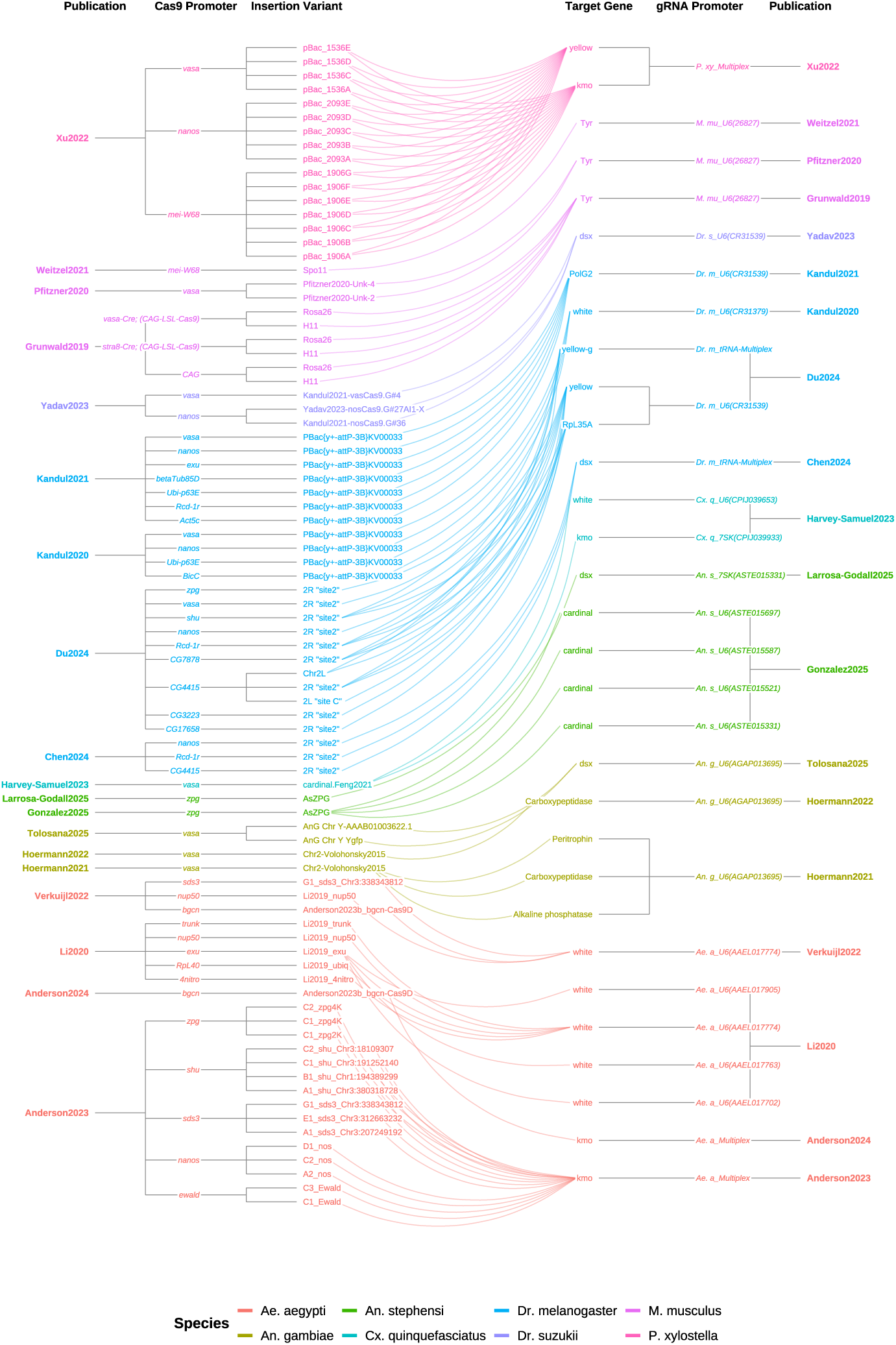
Hierarchical nesting of design factors in split-drive publications. Tanglegram showing the relationships between publications, Cas9 promoters, genomic insertion sites, target genes and gRNA promoter, for non-autonomous (split) drive architectures. Lines connect factor levels that co-occur in at least one cross.

**Figure S16:**
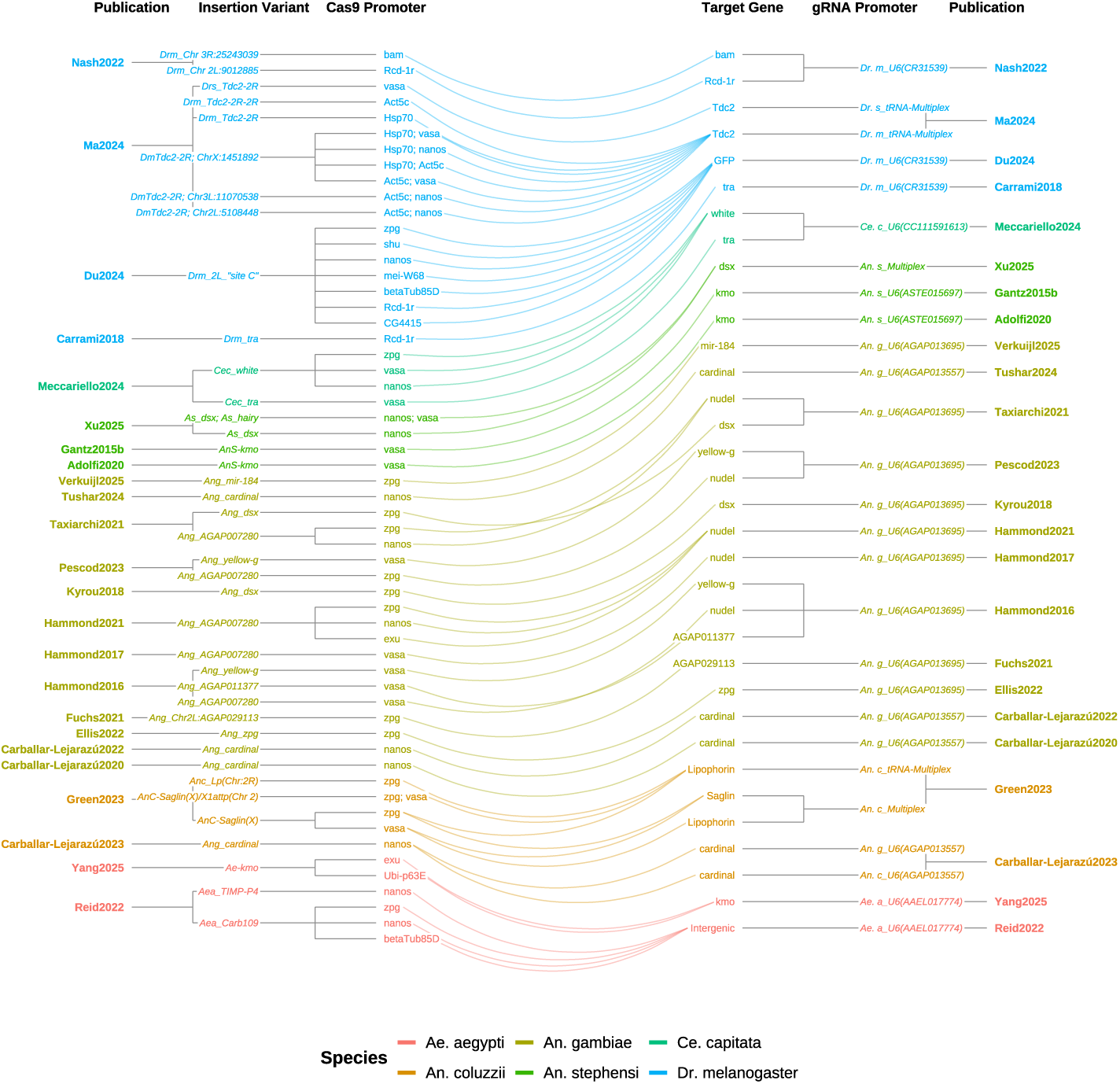
Hierarchical nesting of design factors in autonomous-drive publications. Tanglegram showing the relationships between publications, Cas9 promoters, genomic insertion sites, target genes and gRNA promoter, for autonomous drive architectures. Lines connect factor levels that co-occur in at least one cross.

**Figure S17:**
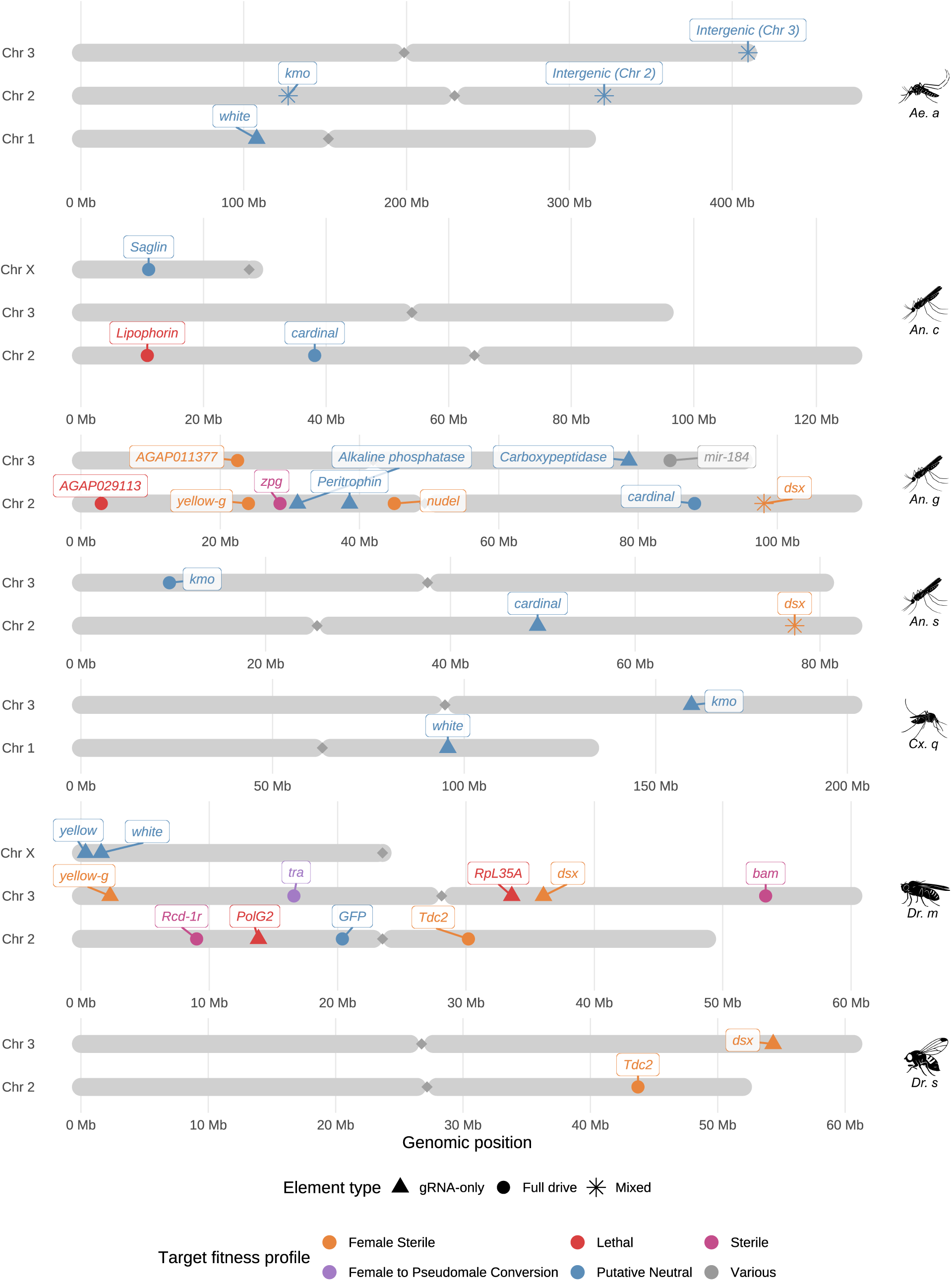
Genomic distribution of gene-drive target loci. For most species, horizontal grey bars depict the chromosomes with separate scaling by species. Labelled markers indicate the genomic coordinates of the drive alleles. Marker shape denotes construct class and marker colour denotes the primary intended fitness/phenotypic effect category. Indents indicate centromere position.

**Figure S18:**
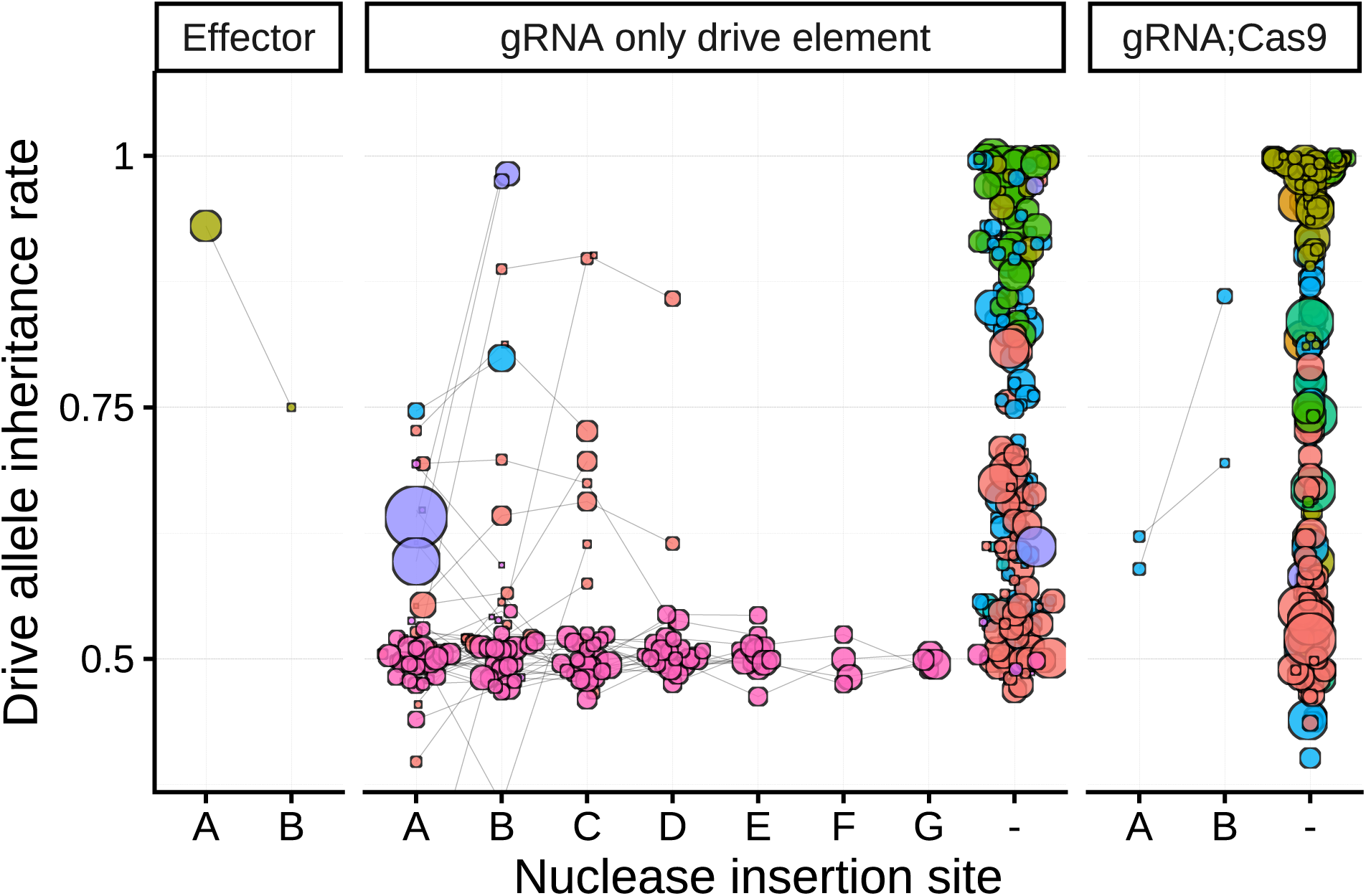
Drive inheritance of Cas9 elements with separate insertion sites. Drive allele inheritance rates for drive constructs paired with a Cas9 element inserted at different genomic locations, with all other design features held constant. This is most commonly achieved through separate transposon mediated insertions of the same Cas9 element. Point size scales with progeny scored. Grey line segments connect crosses equivalent apart from the Cas9 insertion site. ^12;17;22;36;41;46;48;49^ Insertion variants within the same publication receive an arbitrary letter designation; singleton insertions are labelled as “-”.

**Figure S19:**
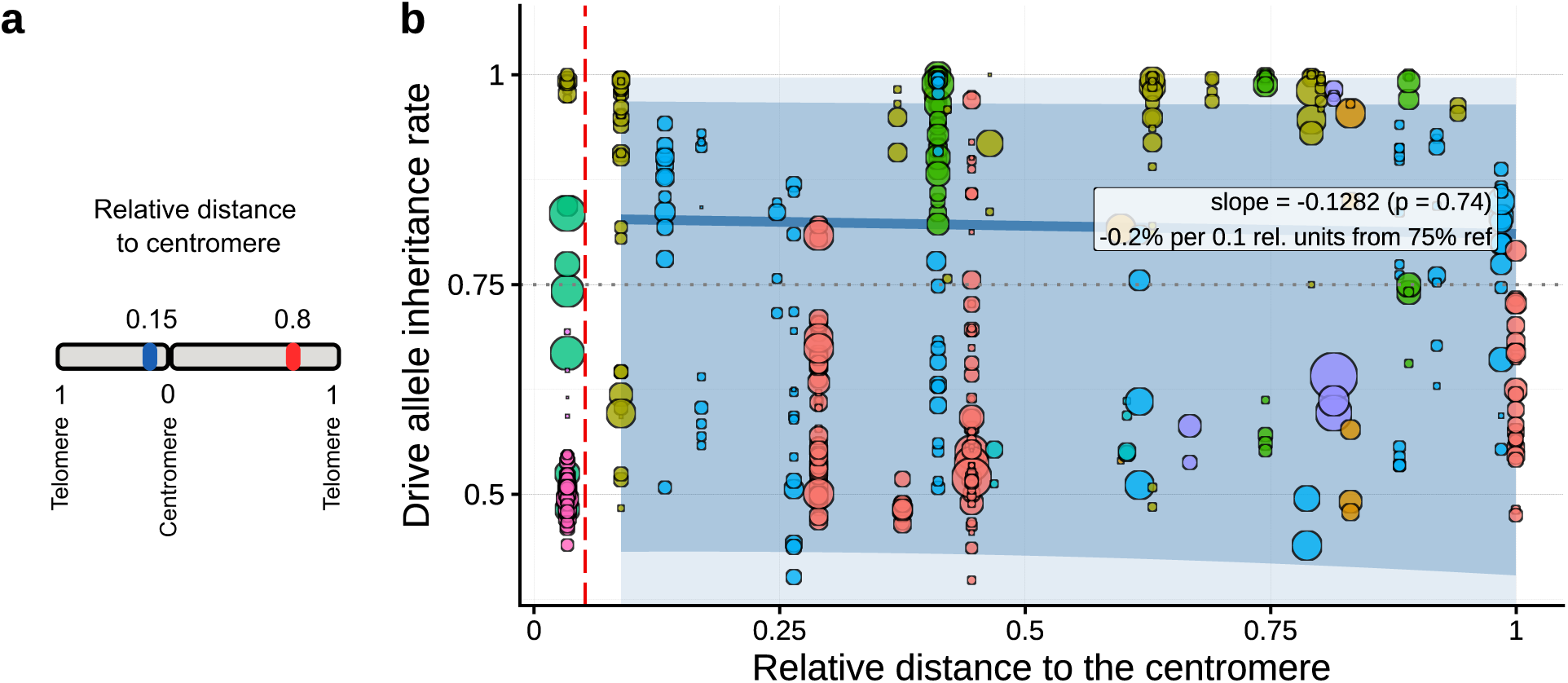
Relationship between centromere proximity and drive inheritance. Drive allele inheritance rates plotted against the relative distance of the drive insertion site from the centromere (normalised by chromosome arm length). Point size scales with progeny scored.

**Figure S20:**
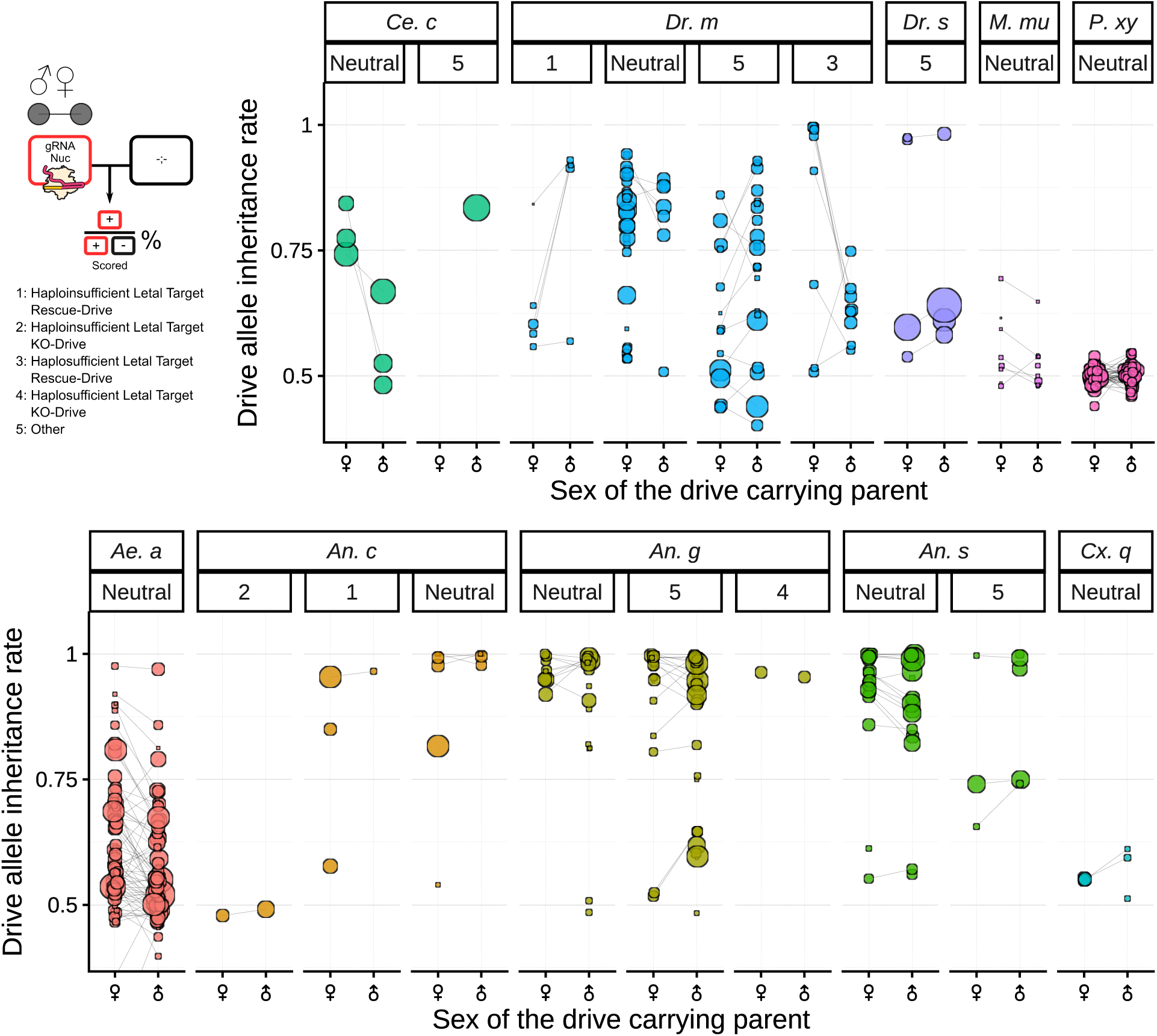
Drive allele inheritance rates by drive-parent sex. Inheritance rates stratified by target-gene fitness profile. “Neutral” indicates putatively neutral targets; numbered lethal-target architectures (inset) distinguish haploinsufficient versus haplosufficient lethal targets crossed with rescue versus knockout drives. Point size scales with progeny scored. Grey line segments connect crosses equivalent apart from drive-parent sex. ^9;10;12–14;16–26;28;32–34;38;40–50^

**Figure S21:**
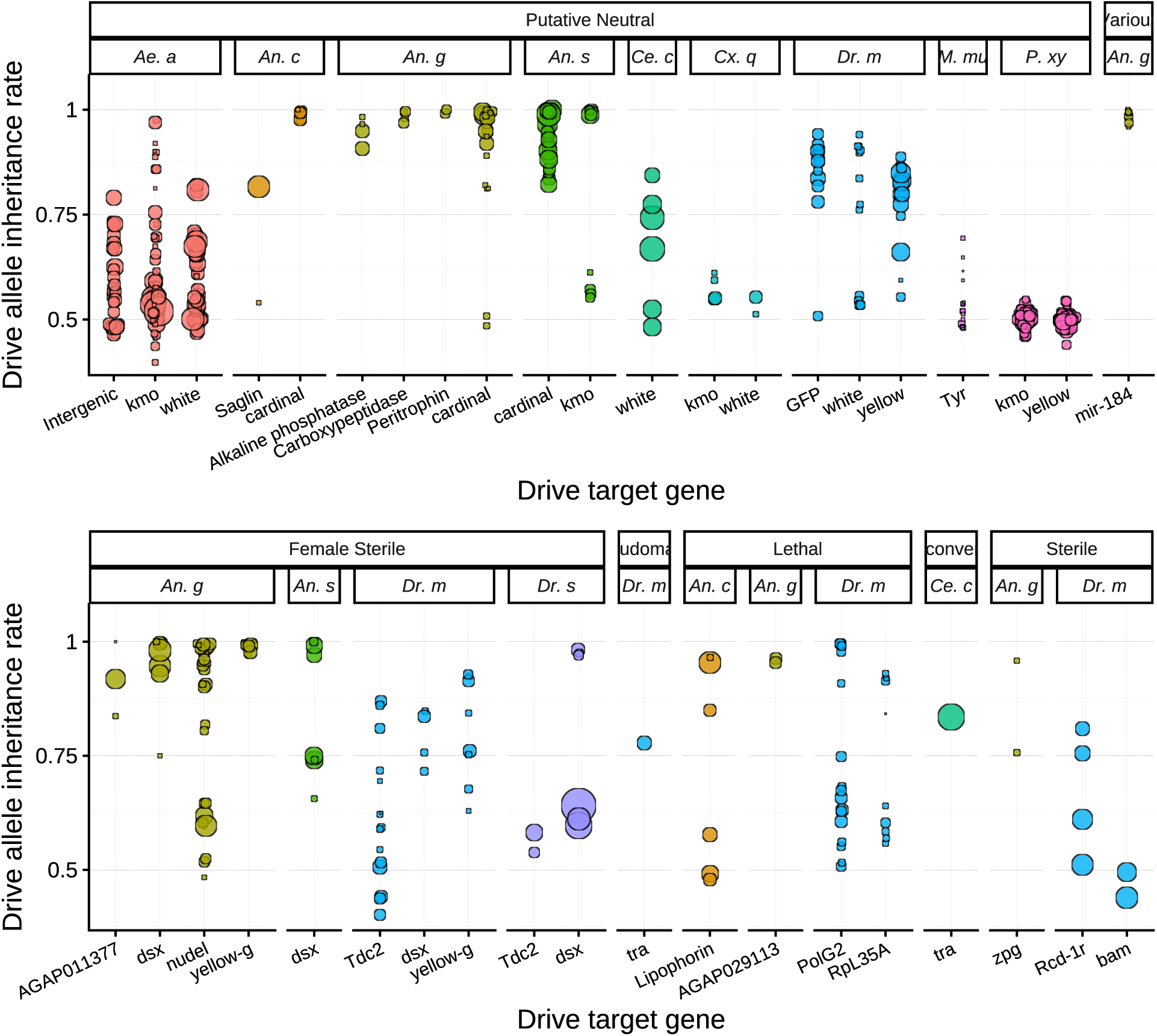
Drive allele inheritance rates stratified by the fitness class of the targeted gene. Observed drive allele inheritance rates (y-axis), grouped by drive target gene (x-axis) and faceted by target-gene fitness classes.

**Figure S22:**
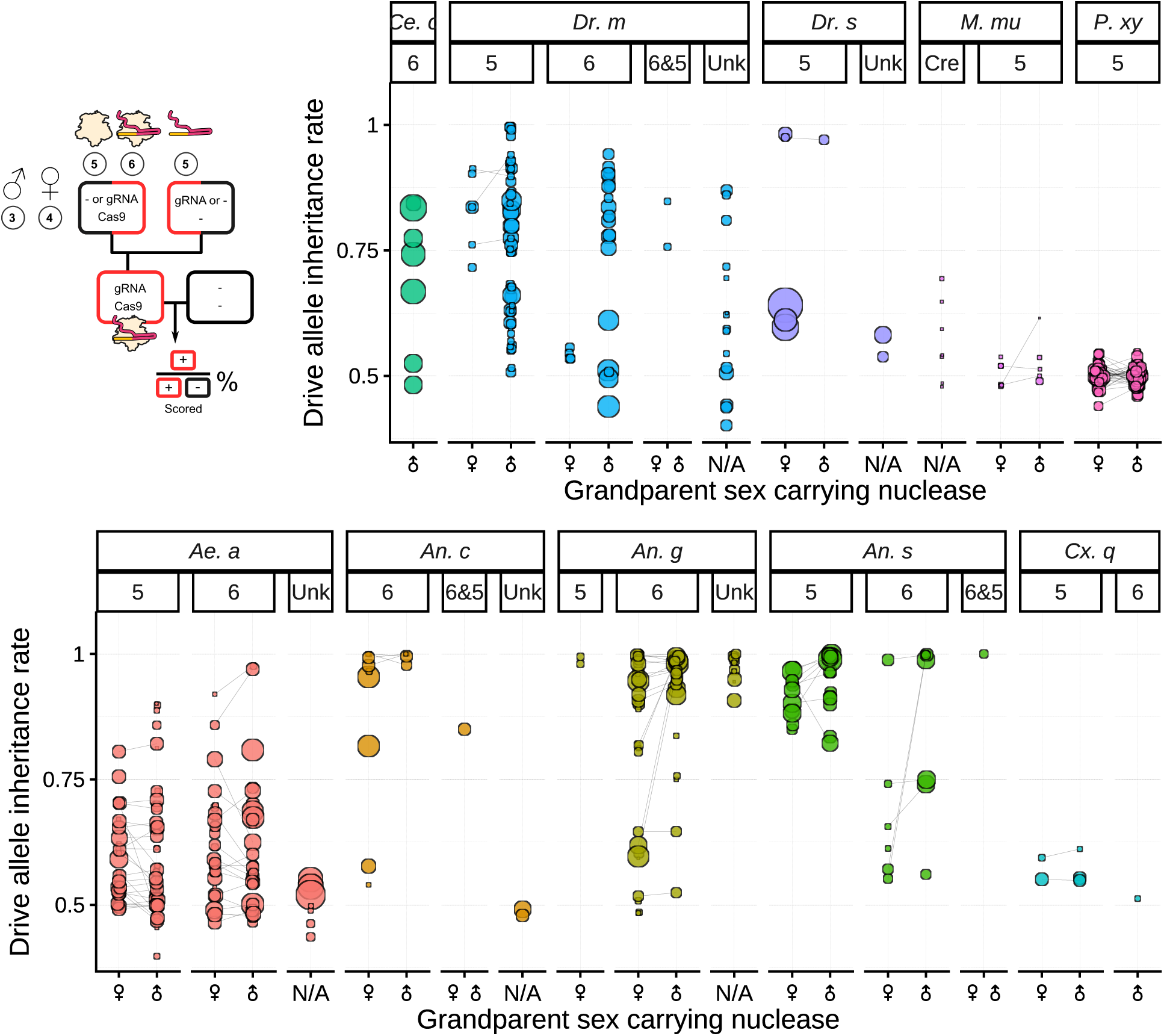
Drive allele inheritance rates by sex of the Cas9-carrying grandparent. Data are faceted by grandparent-cross profile. Facet codes: 5: *−*;Cas9 *×* gRNA;*−*, 6: gRNA;Cas9 *× −*;*−*, 6&5: gRNA;Cas9 *× −*;Cas9, Cre: Crienducible Cas9 (inactive in the grandparent); Unk, unknown. Point size scales with progeny scored. Grey line segments connect crosses equivalent apart from grandparent sex. ^9;14;15;19;21;22;24–26;28;31;34;38;41–47;49^

**Figure S23:**
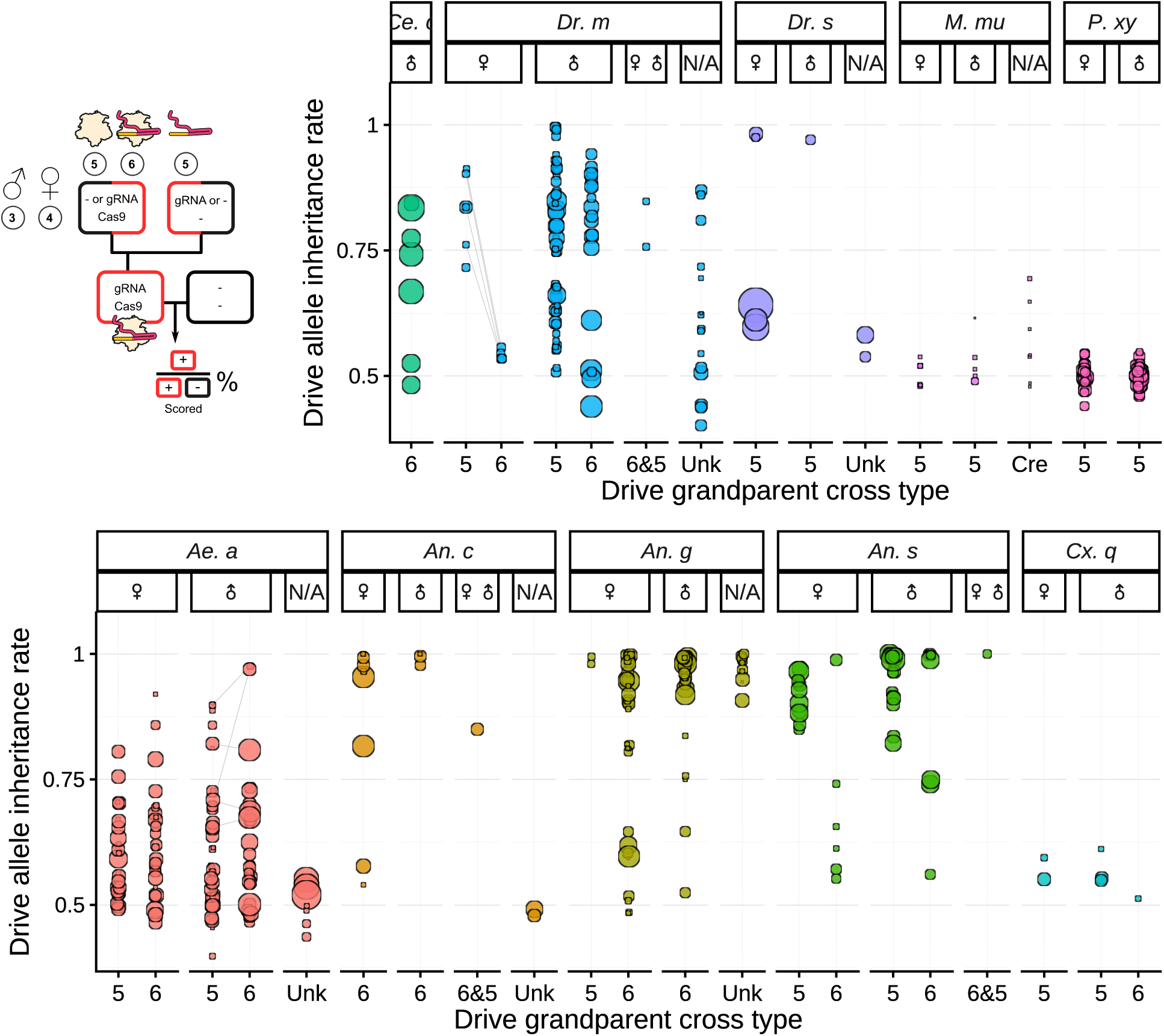
Drive allele inheritance rates by grandparent-cross profile. Data are faceted by the sex of the Cas9-carrying grandparent. Facet codes: 5: *−*;Cas9 *×* gRNA;*−*, 6: gRNA;Cas9 *× −*;*−*, 6&5: gRNA;Cas9 *× −*;Cas9, Cre: Cre-inducible Cas9 (inactive in the grandparent); Unk, unknown. Point size scales with progeny scored. Grey line segments connect crosses equivalent apart from the grandparent-cross profile. ^15;22;24^

**Figure S24:**
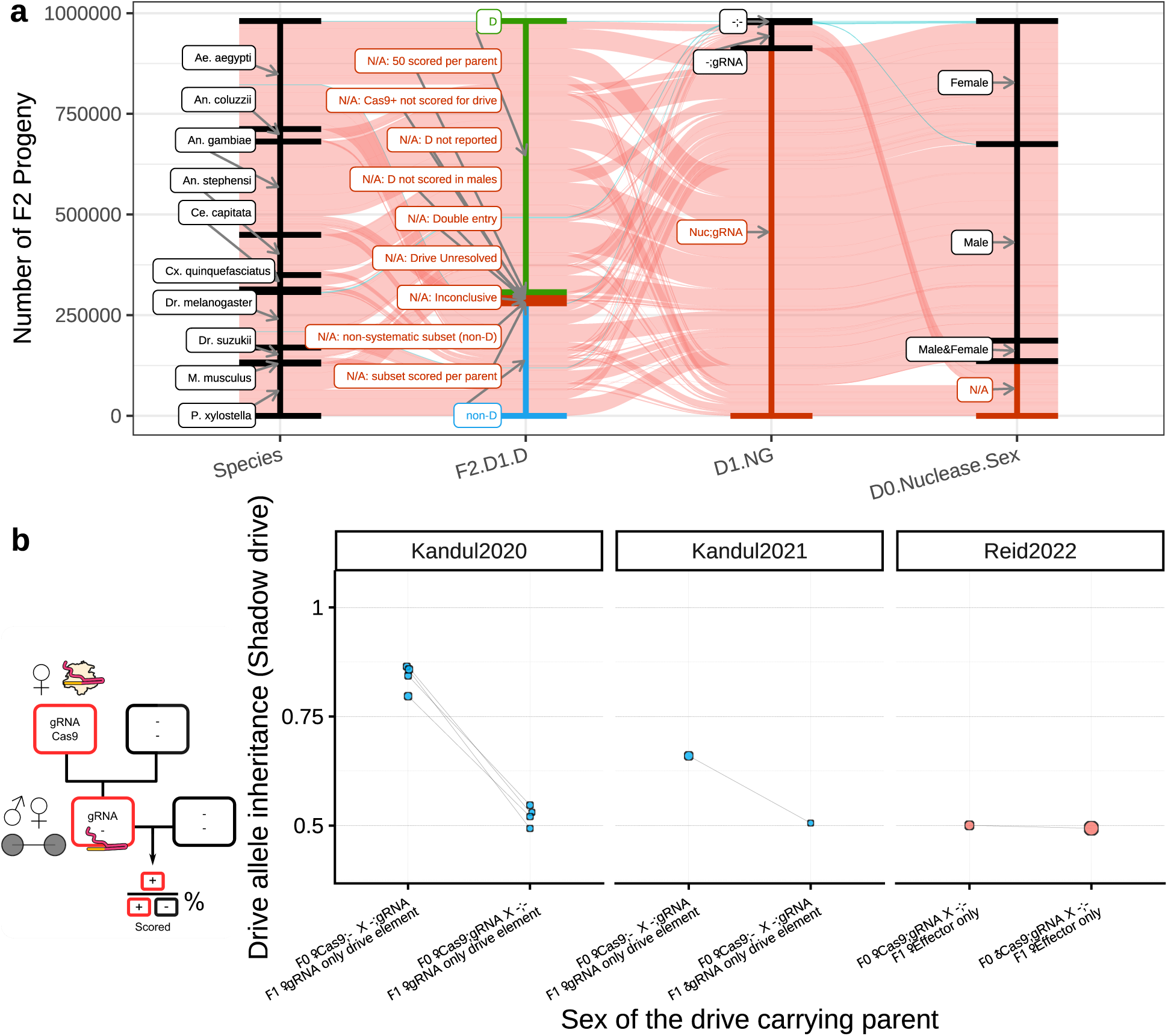
Inheritance bias through deposited Cas9 (shadow drive). **(a)** Data-flow summary showing F2 progeny counts across sequential inclusion criteria, with Cas9 carrying parents excluded. Stream widths are proportional to progeny counts. **(b)** Left: schematic of shadow drive, where drive allele inheritance occurs only through Cas9 deposition. Right: fraction of offspring inheriting the drive from a parent lacking Cas9, faceted by publication. Point size scales with progeny scored. Grey line segments connect crosses equivalent apart from F0 cross type, F1 sex, or F0 Cas9-carrying sex. ^15;16;19^

**Figure S25:**
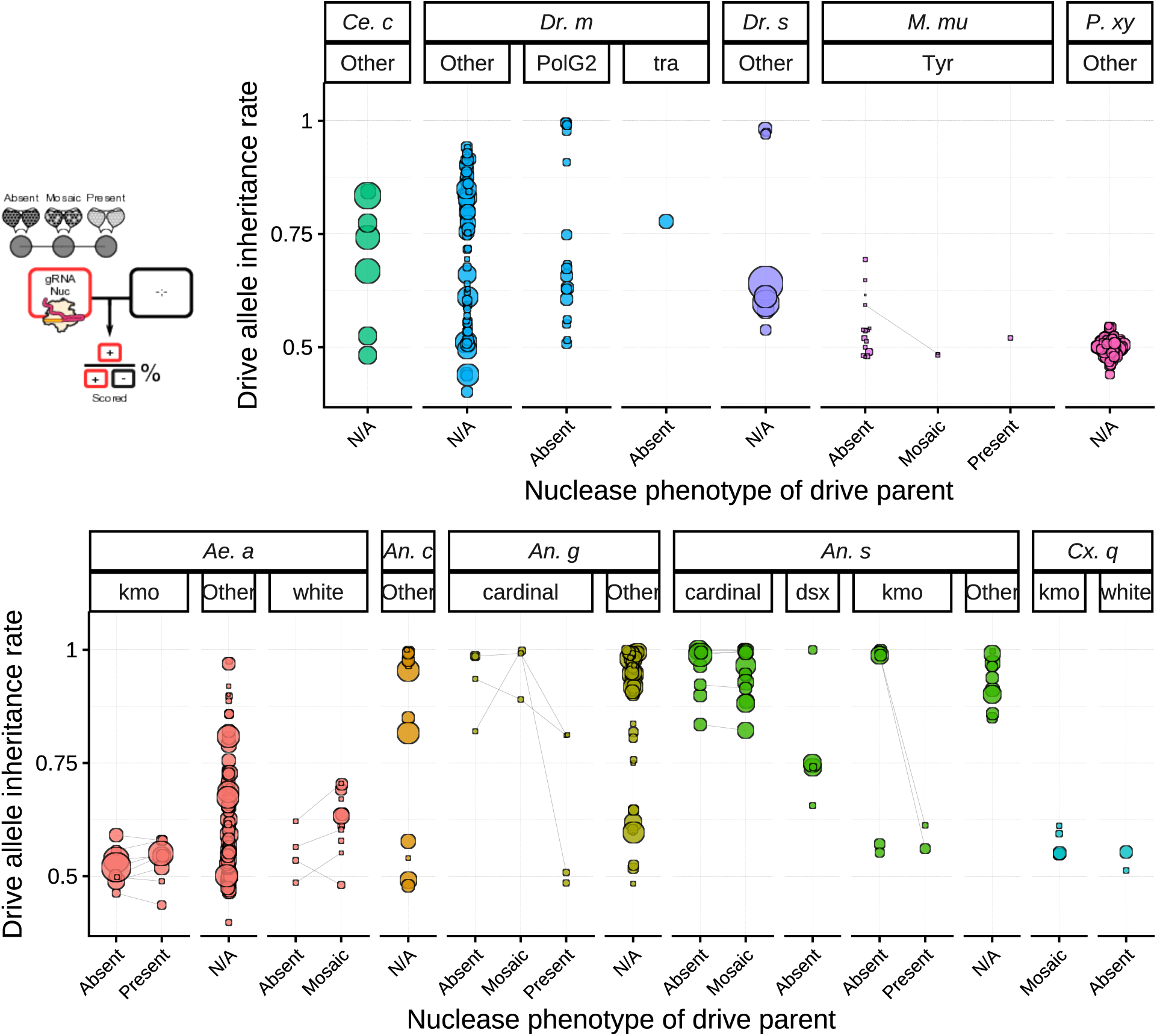
Association between D1 somatic phenotype and drive inheritance. Phenotype categories: Absent (no visible phenotype), Present (knockout phenotype), Mosaic (specifically indicated as mosaic). Panels are faceted by species and target gene; genes without phenotype data are grouped as “Other”. Top-left inset: schematic of phenotypic scoring (WT vs mosaic vs KO). Point size scales with progeny scored. Grey line segments connect crosses equivalent apart from D1 somatic phenotype. ^9;21;25;37;43;48^

**Figure S26:**
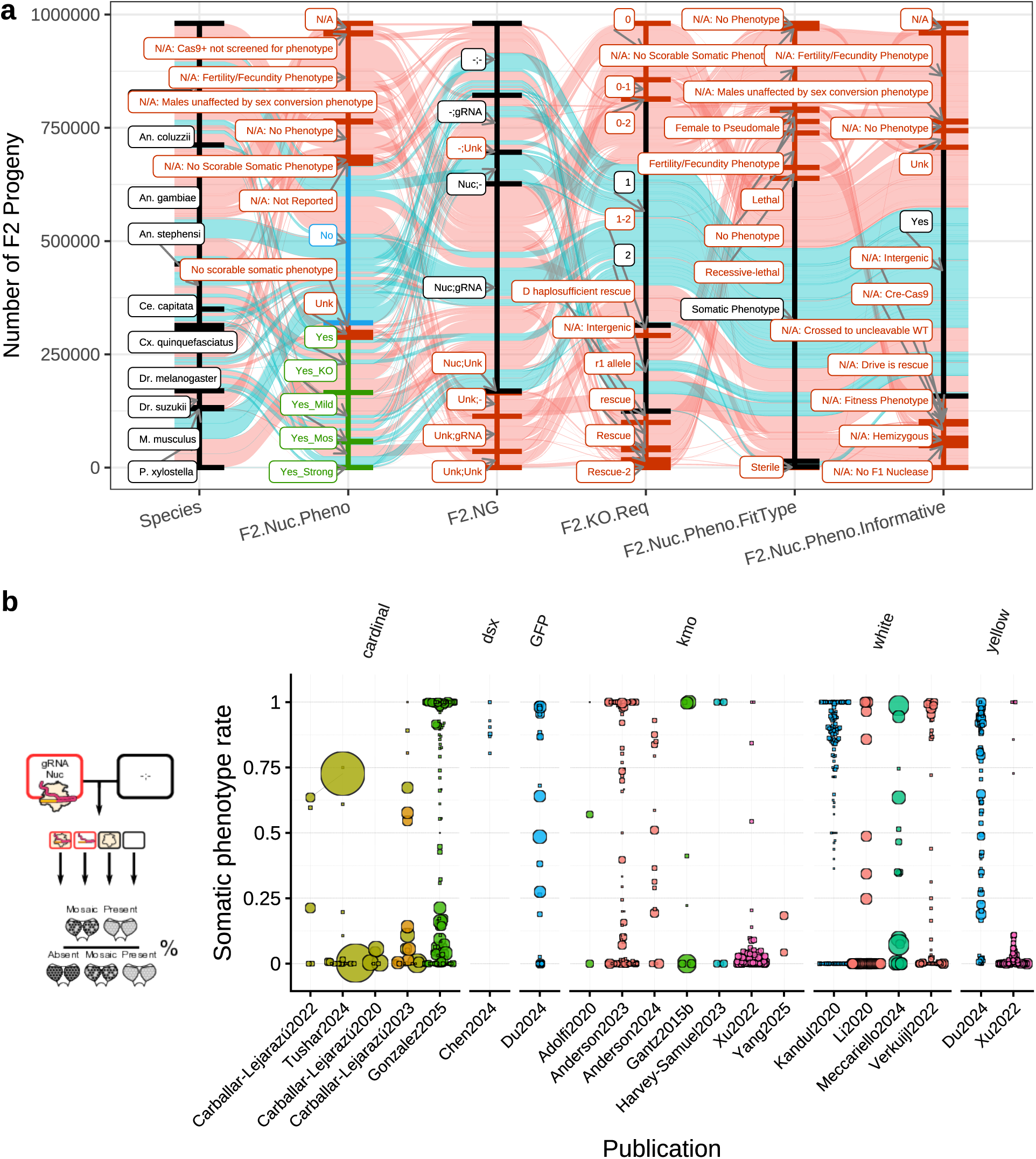
Nuclease-induced somatic phenotype rates across species and target genes. Observed rates of knock-out and mosaic somatic phenotypes in F2 progeny, stratified by species, target gene, and F2 zygotic genotype. Point size scales with progeny scored.

**Figure S27:**
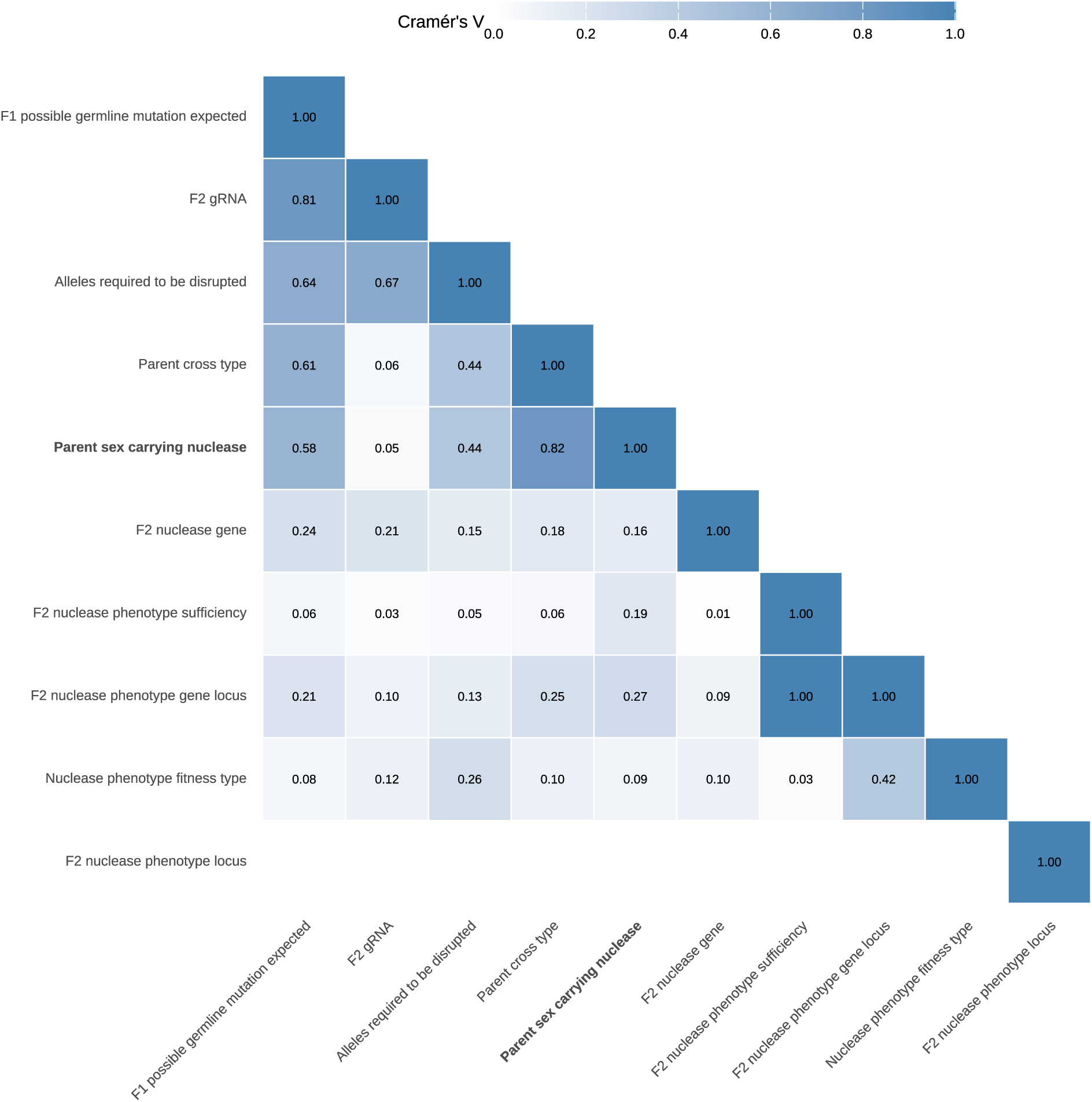
Correlation matrix of F2-level pairing factors for nuclease-induced somatic phenotype. As in the F1 correlation matrix, each cell indicates whether pairs of pairing factors co-vary across the dataset.

**Figure S28:**
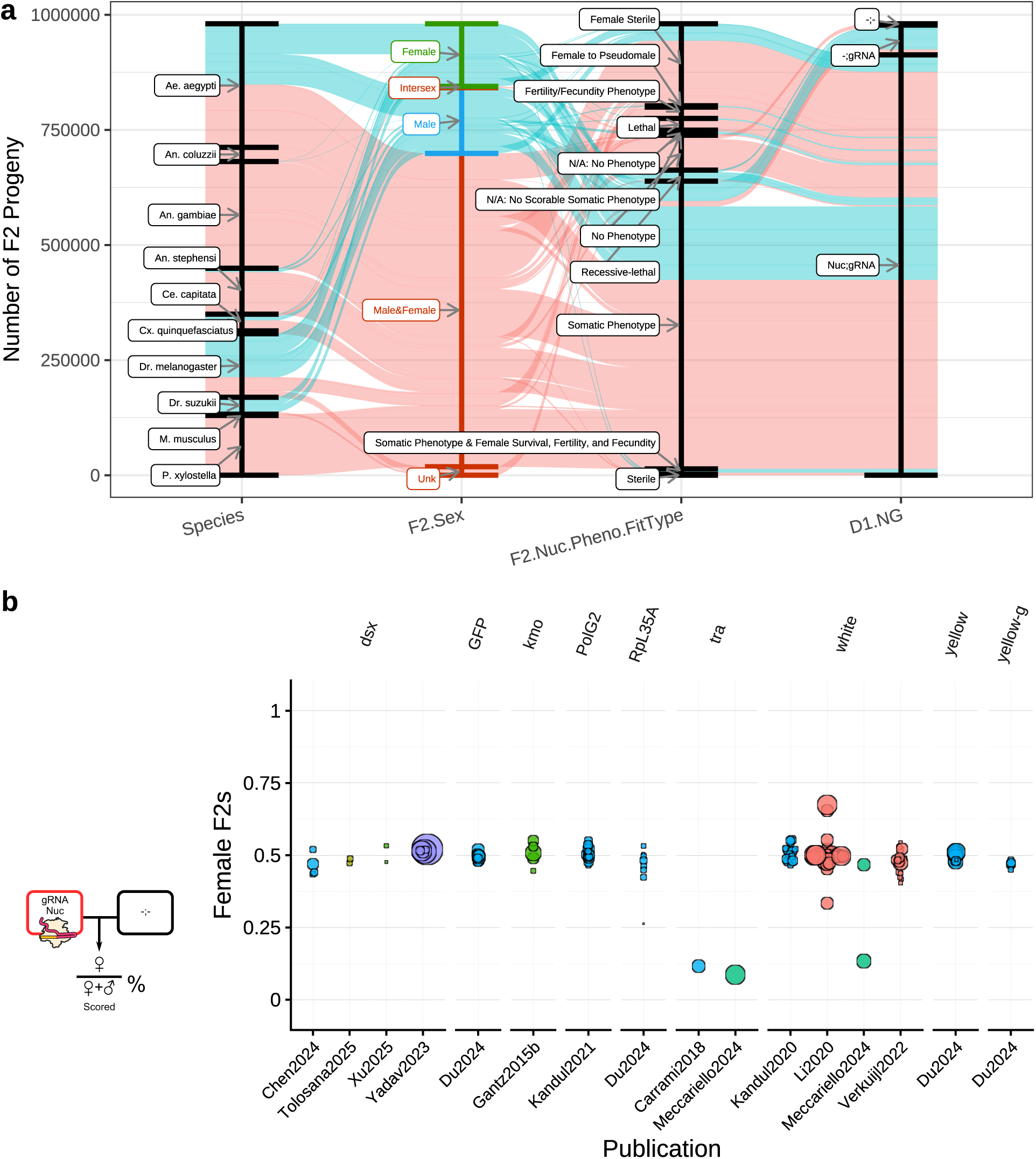
Fraction of female offspring. **(a)** Data-flow summary showing data that are included in F2 sex count. Stream widths are proportional to progeny counts; blue streams have sex recorded, red streams lack sex data. **(b)** Fraction of female F2 progeny per cross, grouped by target gene and publication. Departures reflect sex-linked targets (e.g., *white* in *Ae. aegypti*) or sex-conversion phenotypes (e.g., *tra*). Point size scales with progeny scored.

**Figure S29:**
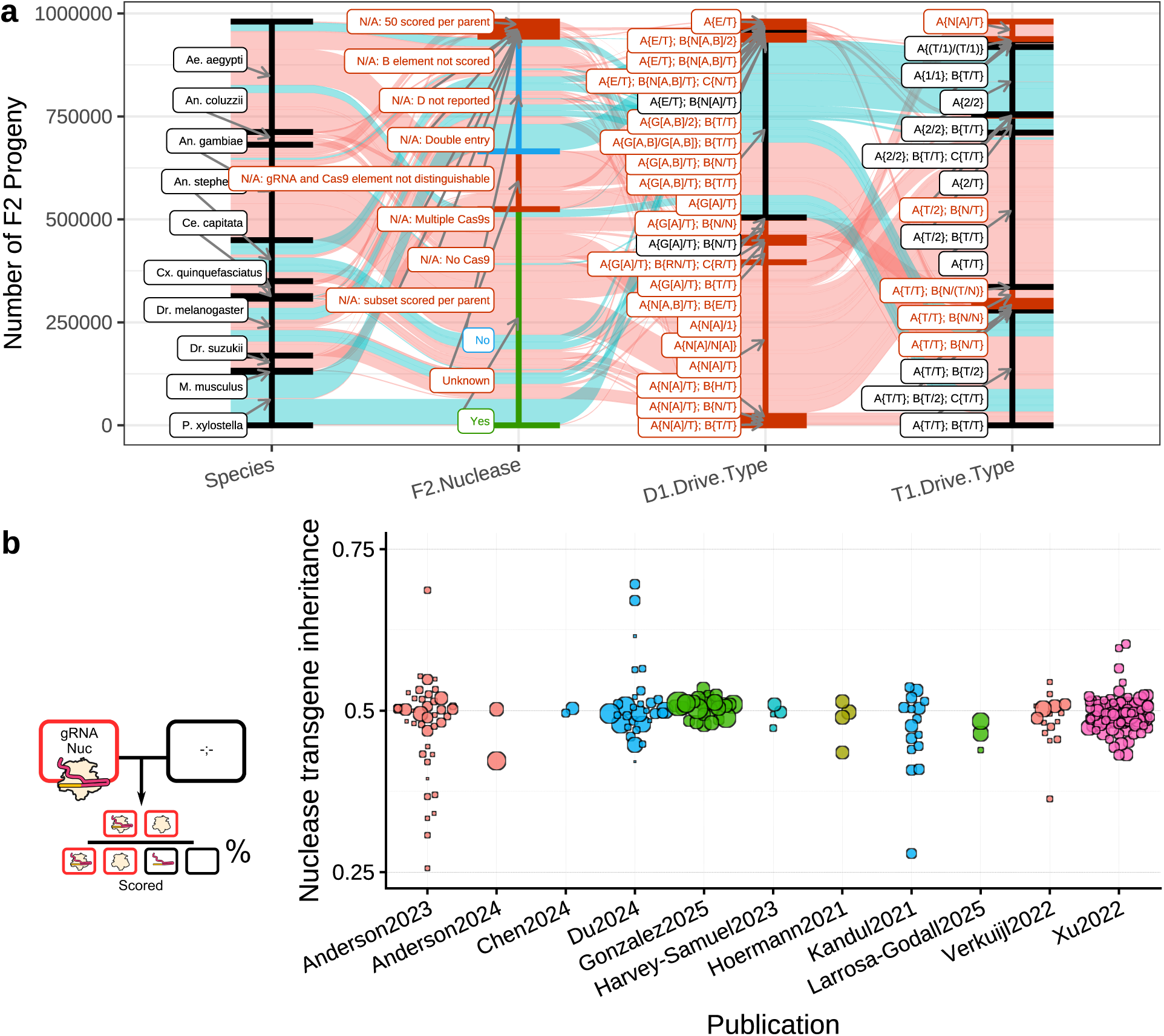
Inheritance of the nuclease transgene in split-drive crosses. **(a)** Data-flow summary for selecting crosses in which the drive element and Cas9 are on separate genetic elements. Stream widths are proportional to F2 progeny counts. **(b)** Fraction of F2 progeny inheriting the nuclease transgene, grouped by publication. Point size scales with progeny scored.

**Figure S30:**
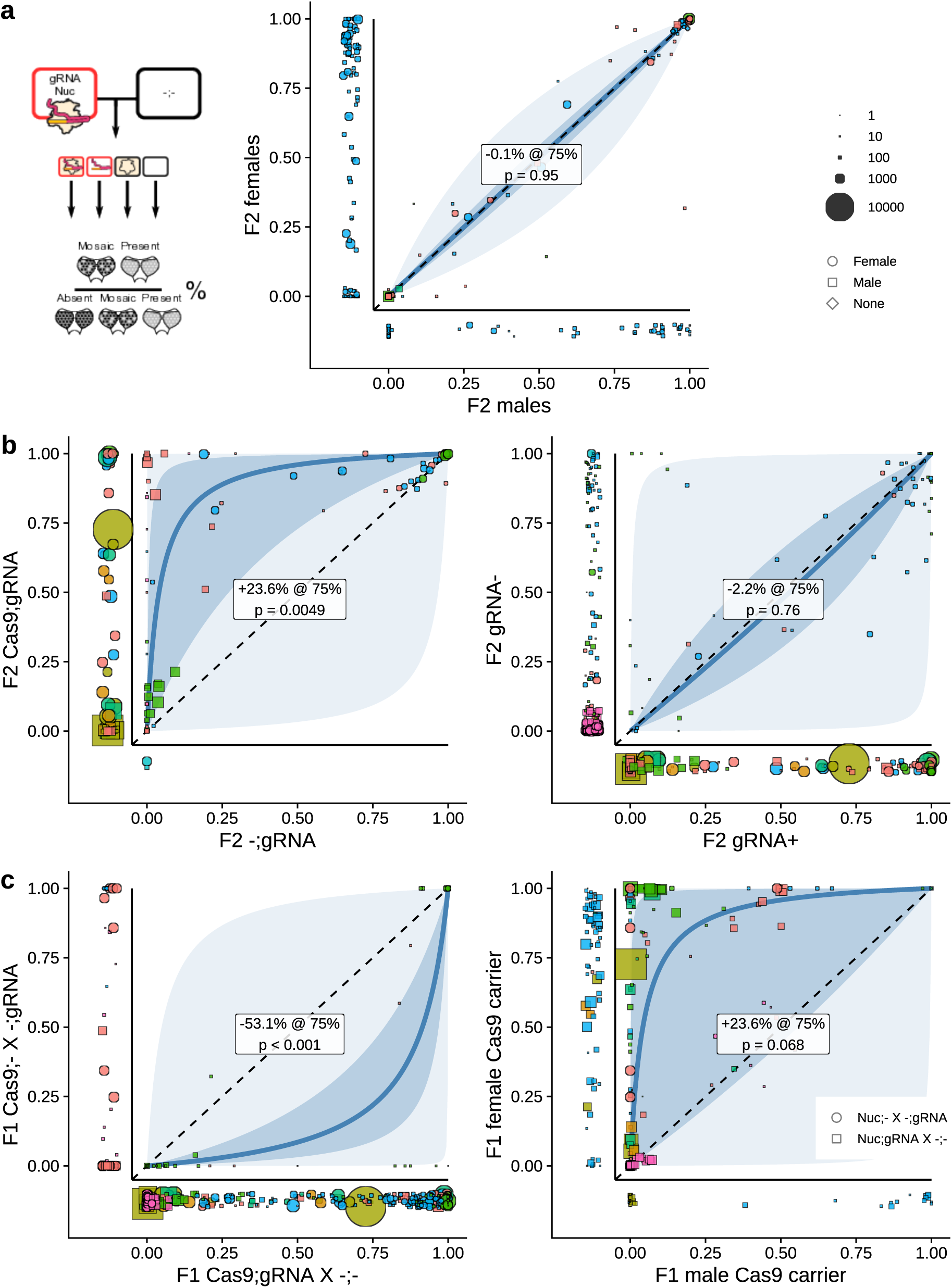
Somatic phenotype rates across F2 and F1 configurations. **(a)** Somatic phenotype rates in female (y-axis) versus male (x-axis) F2 progeny for paired crosses. **(b)** Left: somatic phenotype rates in F2 progeny inheriting both Cas9 and gRNA (y-axis) versus those inheriting gRNA only (x-axis). Right: phenotype rates in F2 progeny carrying the gRNA without a possible germline resistance mutation (x-axis) versus those lacking the gRNA but inheriting a possible germline mutation (y-axis). **(c)** Left: somatic phenotype rates when the Cas9-carrying parent also carried the gRNA (x-axis) versus when Cas9 and gRNA were contributed by different parents (y-axis). Right: phenotype rates when the nuclease-carrying parent was male (x-axis) versus female (y-axis). In all panels, divergence from the diagonal indicates sensitivity to the contrasted factor. Point size scales with progeny scored.

**Figure S31:**
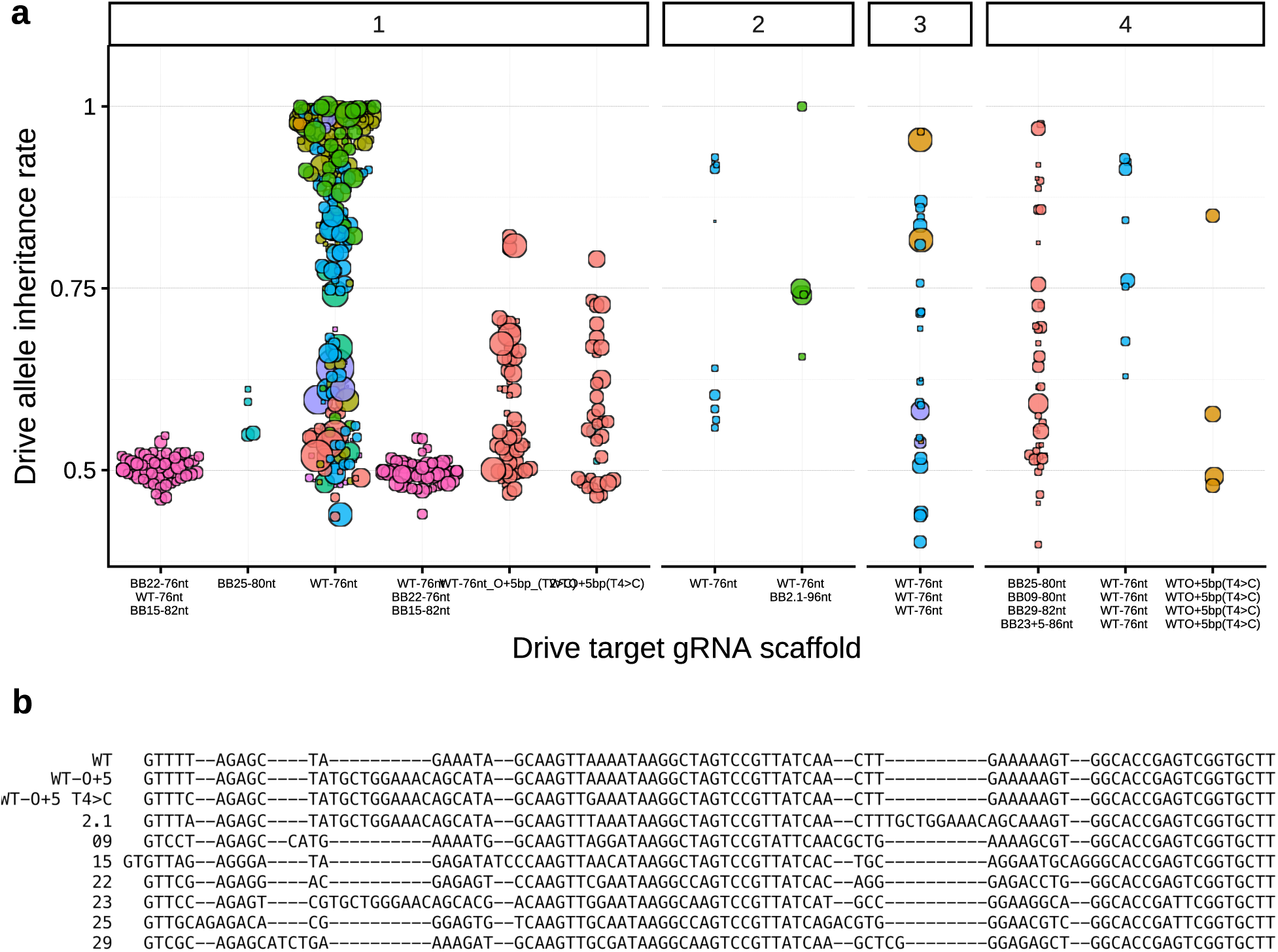
Drive inheritance by gRNA scaffold variant. **(a)** Drive inheritance by gRNA scaffold variant, faceted by unique gRNA count. Modified scaffolds have been adopted primarily to reduce internal sequence homology in multiplexed drive designs. **(b)** Sequence of gRNA scaffold variants.

**Figure S32:**
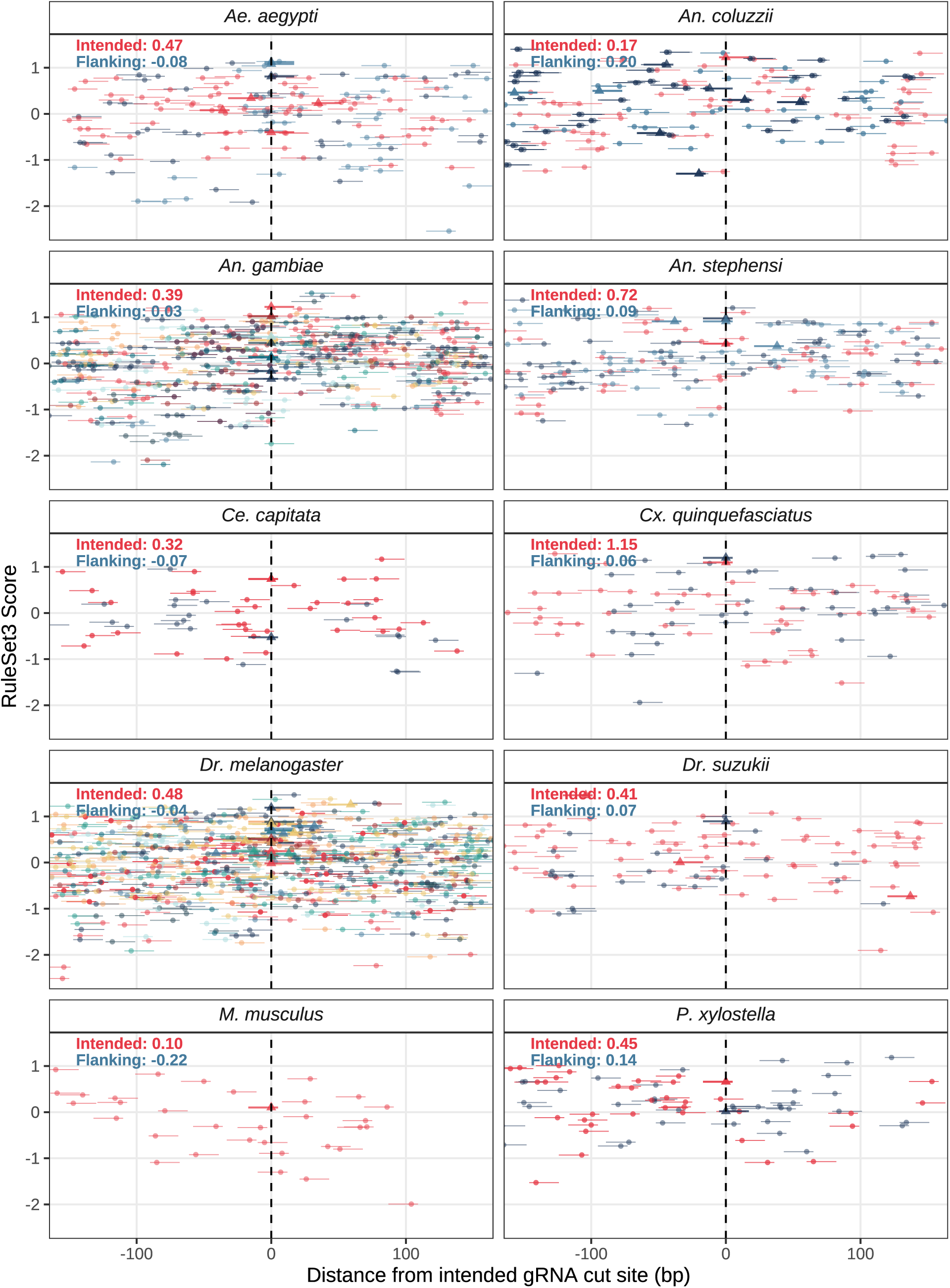
Efficiency scores of flanking gRNAs. RuleSet3 on-target efficiency scores for all candidate SpCas9 gRNAs (NGG-PAM) within 150 bp of each drive’s target site. gRNAs used in published constructs are shown as black triangles; flanking candidates as circles. Horizontal segments indicate spacer and PAM footprints. Panels are faceted by species; annotated values summarise scores for intended versus flanking gRNAs. We did not filter flanking gRNAs by off-target matches.

**Figure S33:**
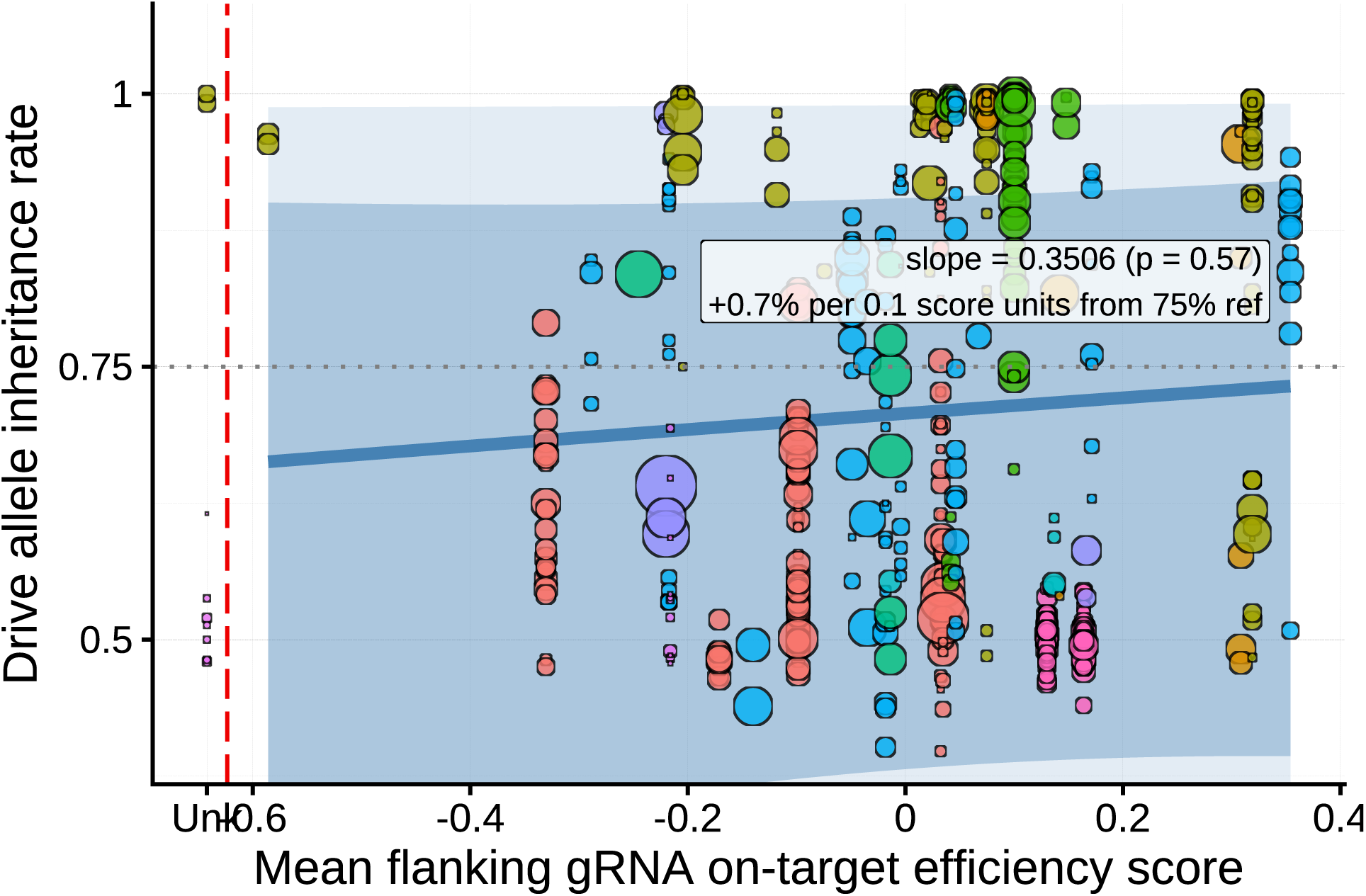
Mean flanking gRNA efficiency score versus drive inheritance. Drive allele inheritance rates plotted against the mean RuleSet3 efficiency score of gRNAs flanking each drive’s target site. Point size scales with progeny scored.

**Figure S34:**
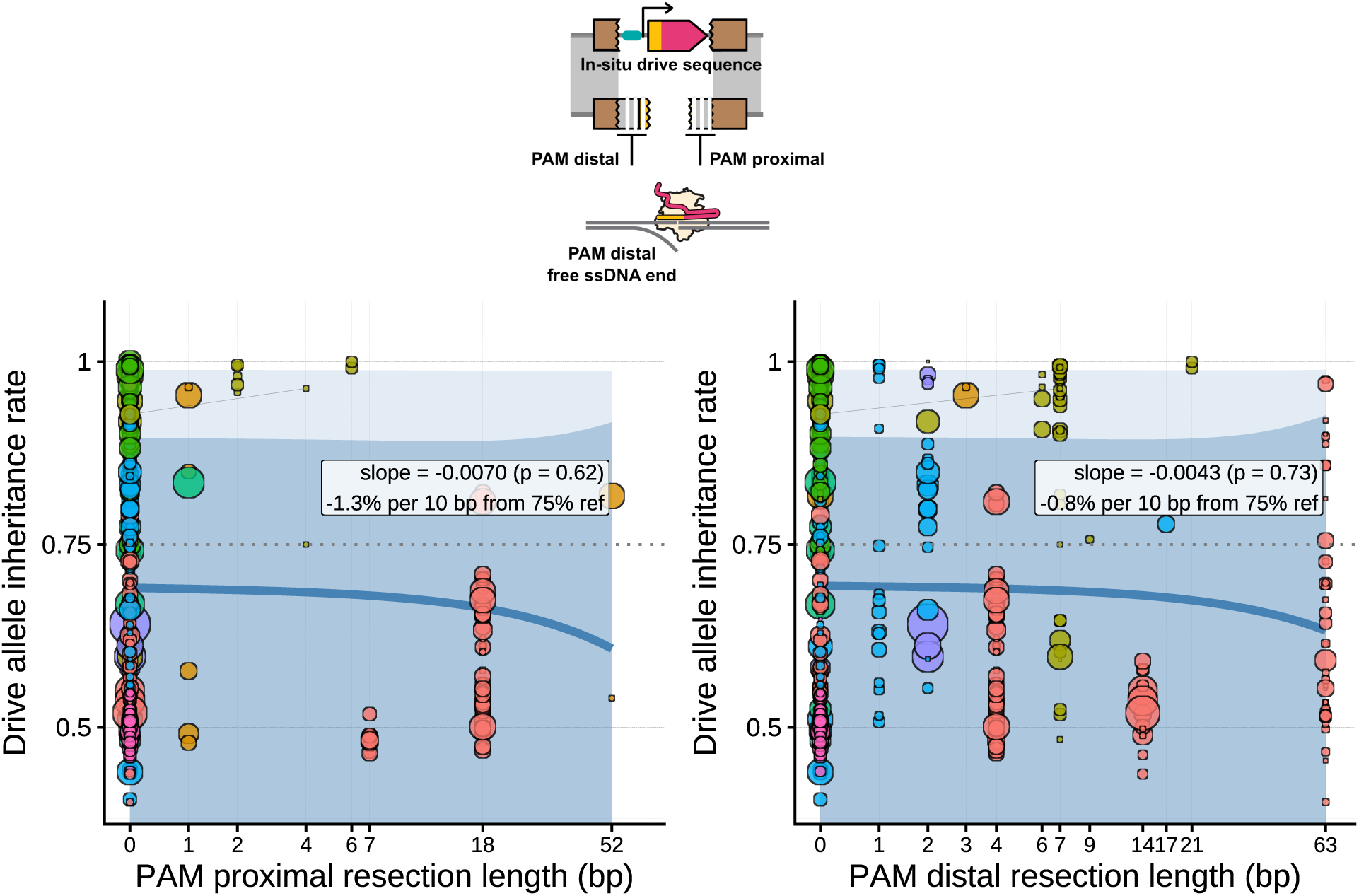
Resection distance by PAM orientation. Drive allele inheritance rates plotted against the non-alignment length on each side of the gRNA cut site, separated by whether the side is PAM-proximal or PAM-distal. The asymmetry of Cas9 cleavage relative to the PAM may produce unequal resection requirements on each side of the break. Point size scales with progeny scored. ^36^

**Figure S35:**
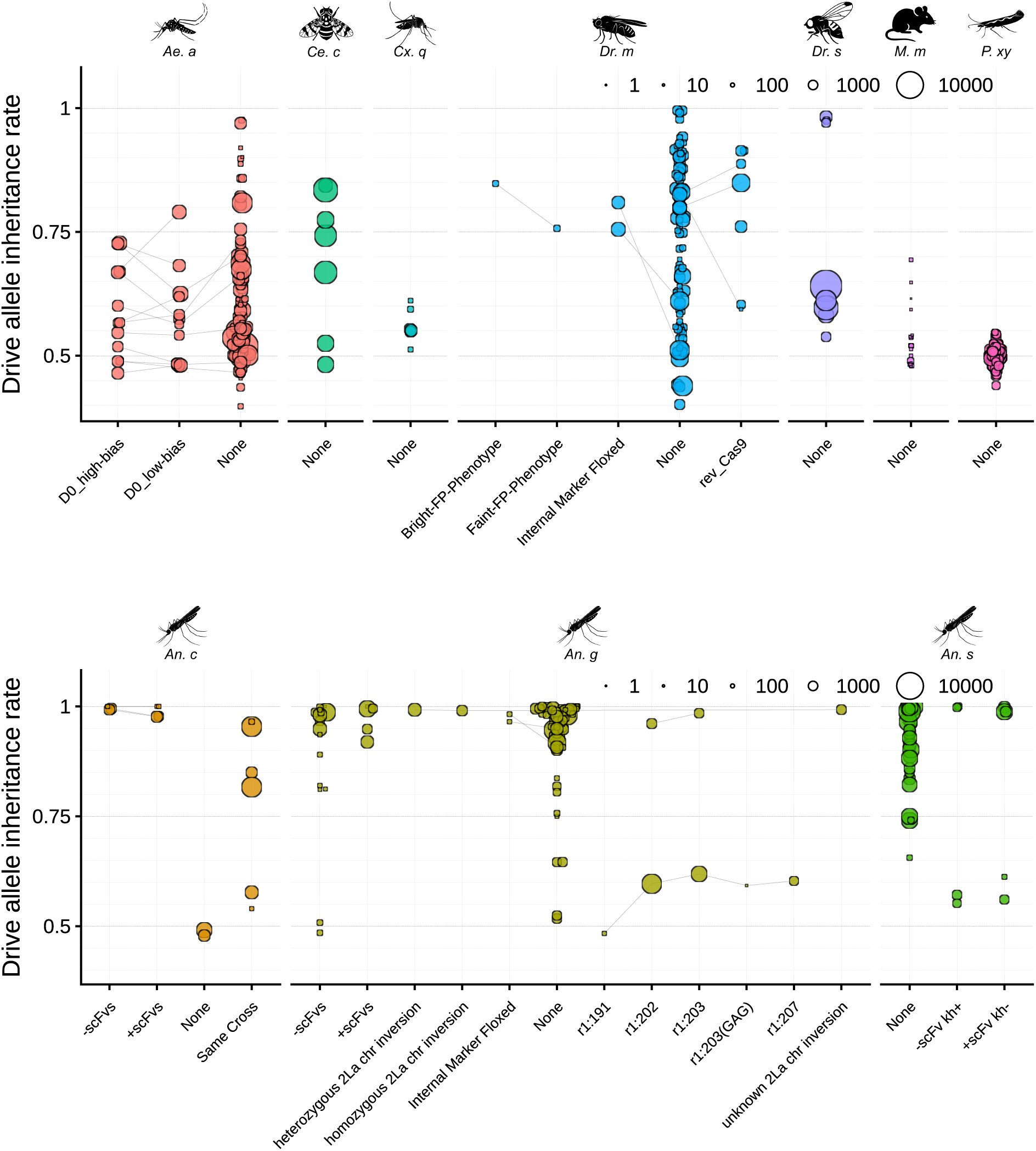
Drive allele inheritance stratified by miscellaneous cross-level factors. Inheritance rates are shown against categories of idiosyncratic study-specific differences that could not be mapped to the primary moderators. Grey line segments connect crosses equivalent apart from the miscellaneous factor. ^11;12;19;28;31;32;35;40^ “NA” denotes crosses with no recorded miscellaneous factor. By default, crosses differing in these factors are visualised separately in other figures; the miscellaneous factor is substituted when another moderator captures the same variation (e.g., drive size varies with marker and cargo presence). Point size scales with progeny scored.

**Figure S36:**
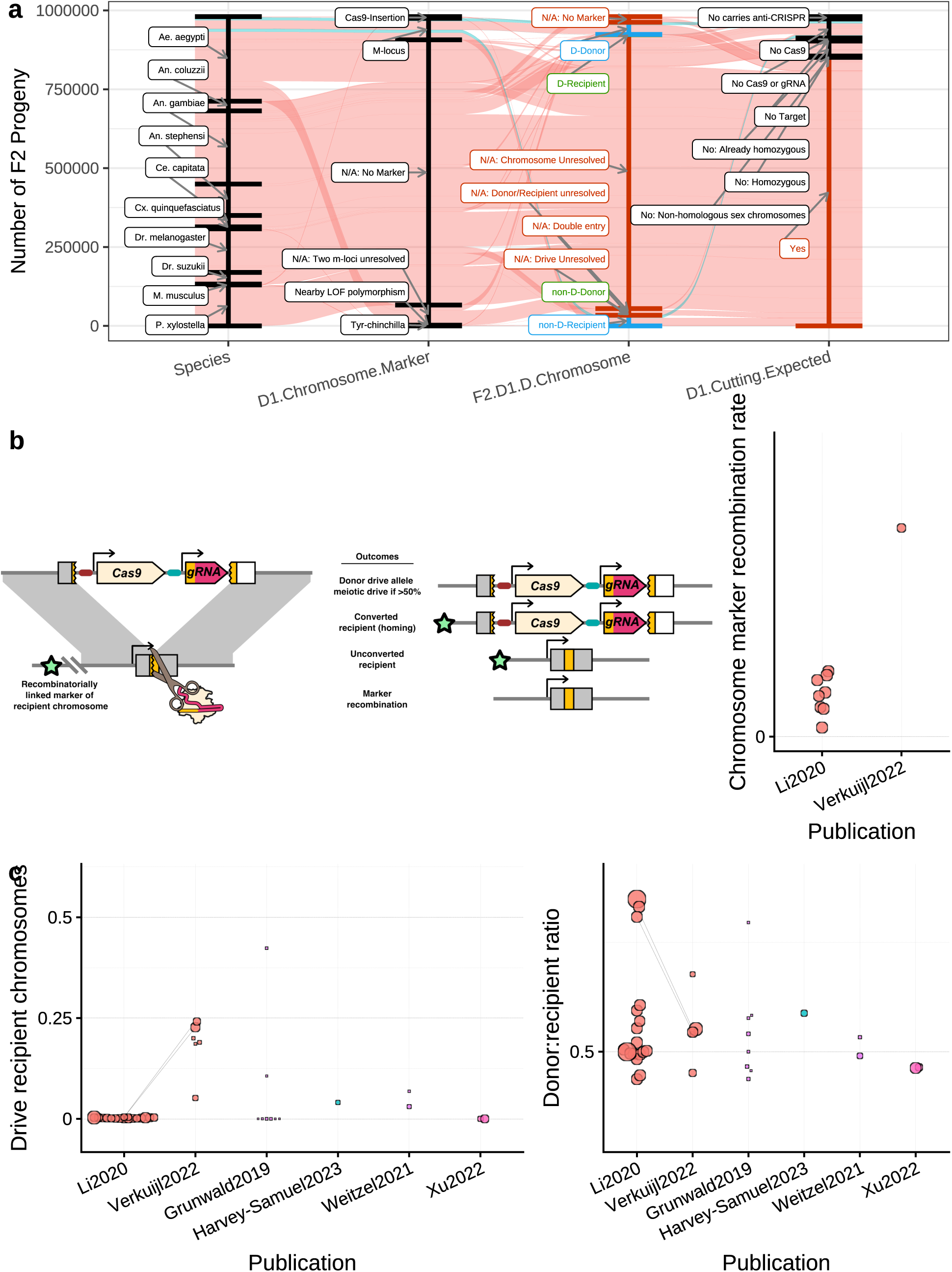
Recombinatorily linked markers differentiate homing from meiotic drive. **(a)** Data-flow summary of F2 progeny counts across species, marker type, and allele classification (drive-carrying, non-drive, donor, recipient chromosome). Background recombination uses non-cutting crosses only; homing and meiotic drive use cutting crosses. **(b)** Schematic of the inference logic when the drive allele is tracked via a linked marker on the recipient chromosome. Graph shows background recombination rates from control crosses; low rates are required for recombination to serve as a homing indicator. **(c)** Left: fraction of recipient chromosomes carrying the drive allele (inferred homing). Right: donor versus recipient chromosome inheritance (inferred meiotic drive). Point size scales with progeny scored. Grey line segments connect crosses equivalent apart from the publication of origin. D1 somatic phenotype is excluded from pairing in this figure to link otherwise equivalent crosses across studies.

